# FRAP-in-SR: Fluorescence recovery in the Super-Resolution regime reveals subcompartments of 53BP1 foci

**DOI:** 10.1101/2025.05.07.652606

**Authors:** Chengchen Wu, Janeth Catalina Manjarrez-González, Muntaqa Choudhury, Noor Shamkhi, Siwen Ding, Vishnu M Nair, Viji M. Draviam

## Abstract

We combine Lattice Structured Illumination Microscopy (*di*SIM or SIM^2^ with ∼60 nm resolution), Lattice Light-sheet microscopy and Fluorescence Recovery After Photobleaching (FRAP) to explore 53BP1 dynamics in Retinal Pigment Epithelial cells. 53BP1 forms liquid condensates during double-strand DNA repair, long-range DNA end-joining and heterochromatin maintenance. Our super-resolution movies reveal differences in 53BP1 foci contour: some foci are compact and stationary while others appear amorphous, dynamically changing shapes. To explore them, we developed FRAP in the Super-Resolution regime (FRAP-SR). 53BP1 foci with an amorphous loose contour display subcompartments that recover 53BP1-eGFP signals rapidly, indicating differential protein mobilities and 53BP1 functions within a single foci. In contrast, 53BP1-eGFP foci with a compact contour recover uniformly as single foci but show higher heterogeneity in 53BP1-eGFP recovery rates compared to foci that recover as multiple subcompartments. In cells released from aphidicolin, amorphous foci show faster 53BP1 recovery compared to compact foci. We discuss the conceptual implications of different 53BP1 mobilities, and how the FRAP-SR method transforms studies of dynamic 60-100 nm structures.

## Introduction

Double-strand breaks (DSBs) are highly toxic lesions that, if uncorrected, can cause mutations and chromosomal instability, leading to cancers^1^. DSBs signal a histone modification cascade recognised by dimers of 53BP1 (Tumour Protein p53 Binding protein-1; TP53BP1) leading to the formation of higher-order 53BP1 oligomers and a mature foci structure^2–5^. 53BP1 is a large 1972 a.a long protein forming discrete nuclear foci that enrich downstream checkpoint effectors^6,7^ within a minute of DNA damage. These foci can resolve within two minutes^8^. Similar rapid recruitment and release has been observed in other components of the DNA damage response (DDR) pathway^9^, highlighting 53BP1’s role in a highly dynamic and macromolecular process.

Phase separation of 53BP1 determines the liquid-like behaviour of DNA repair compartments^10^. Beyond DNA repair, 53BP1 regulates heterochromatin structure through phase separation^11^ and facilitates long-range DNA end-joining interactions^12,13^ during VDJ recombination and class switch recombination. 53BP1 foci can also arise in response to replication stress during the G1 phase following mitosis^14,15^. Importantly, 53BP1 foci can remain unresolved for days in multinucleated G1 cells, while laser damage-induced 53BP1 foci within the same nuclei can resolve within minutes^8^. Thus, 53BP1 regulates many macromolecular nuclear events and 53BP1 foci can be stable or dynamic within the same cell^8^.

Various approaches exist to induce DNA damage and track it live using Super-Resolution (SR) microscopy^16^ - SR imaging has revealed the rearrangement of 53BP1 signals during DNA repair^17,18,19^. However, 53BP1 protein dynamics have not been correlated *per se* with the foci’s architectural changes as this requires a combination of FRAP and SR imaging. Single-molecule recovery FRAP, an SR imaging technique used for nuclear envelope (40 nm dimensions), has significantly expanded our understanding of nuclear wall proteins^20,21^, but it requires continuous exposure for prolonged periods to collect signals, making it unsuitable for studying DNA repair, which is highly photosensitive. As opposed to this, Lattice Structured Illumination Microscopy (SIM) is a gentle SR approach and can computationally improve the diffraction-limited lateral resolution by at least two-fold^22,23^. Thus, combining FRAP with Lattice SIM may help probe 53BP1 protein dynamics in addition to simultaneous visualisation of the foci’s architectural changes in the SR regime, but this has not been reported so far.

Using FRAP to capture subcellular changes in the SR regime, we analyse 53BP1 protein dynamics within nuclear structures at 60 nm resolution for the first time. With this new approach, we reveal subcompartments within 53BP1 foci; these subcompartments display faster 53BP1 protein mobility than the others without subcompartments. Based on super-resolved foci contour and FRAP recovery, we find at least two different types of 53BP1 foci: (i) foci that remain compact and recover as a single compartment during the 3-minute FRAP period, but show high heterogeneity in 53BP1-eGFP recovery rates and (ii) foci that show multiple subcompartments during FRAP recovery and display an amorphous contour that is irregular and dynamically shape-changing. While the compact foci appear dormant and largely stationary, the amorphous foci are loose and mobile. Using lattice lightsheet movies of aphidicolin-treated cells, we confirm faster recovery of 53BP1 in amorphous foci compared to compact foci. Thus, by exploiting FRAP-SR as a gentle method to correlate protein diffusion with subcellular structural changes in the SR regime, we show evidence for subcompartments within 53BP1 foci and characterise two distinct 53BP1 foci that differ in their activities.

## Results

### FRAP in the SR regime reveals subcompartments within 53BP1 foci

To first demonstrate that we can achieve lateral resolutions of ∼60 nm in our FRAP-SR imaging setup, we imaged 60 nm DNA origami beads and tested two reconstruction algorithms, SIM or dual iterative SIM (*di*SIM, also called SIM^2^), to measure peak-to-peak distances (Figure 1A and 1B). The twin pattern of origami beads separated by 60 nm is fully resolved (Figure 1B) in *di*SIM but not in SIM-processed images. Measuring the distance between the peak intensities of the twin beads showed a median length of 64.4 nm in 80% of the beads (Supplementary Figure 1A). We conclude that 60 nm lateral resolution can be achieved using the *di*SIM algorithm for image reconstruction in our FRAP-SR imaging setup.

**Figure 1:**
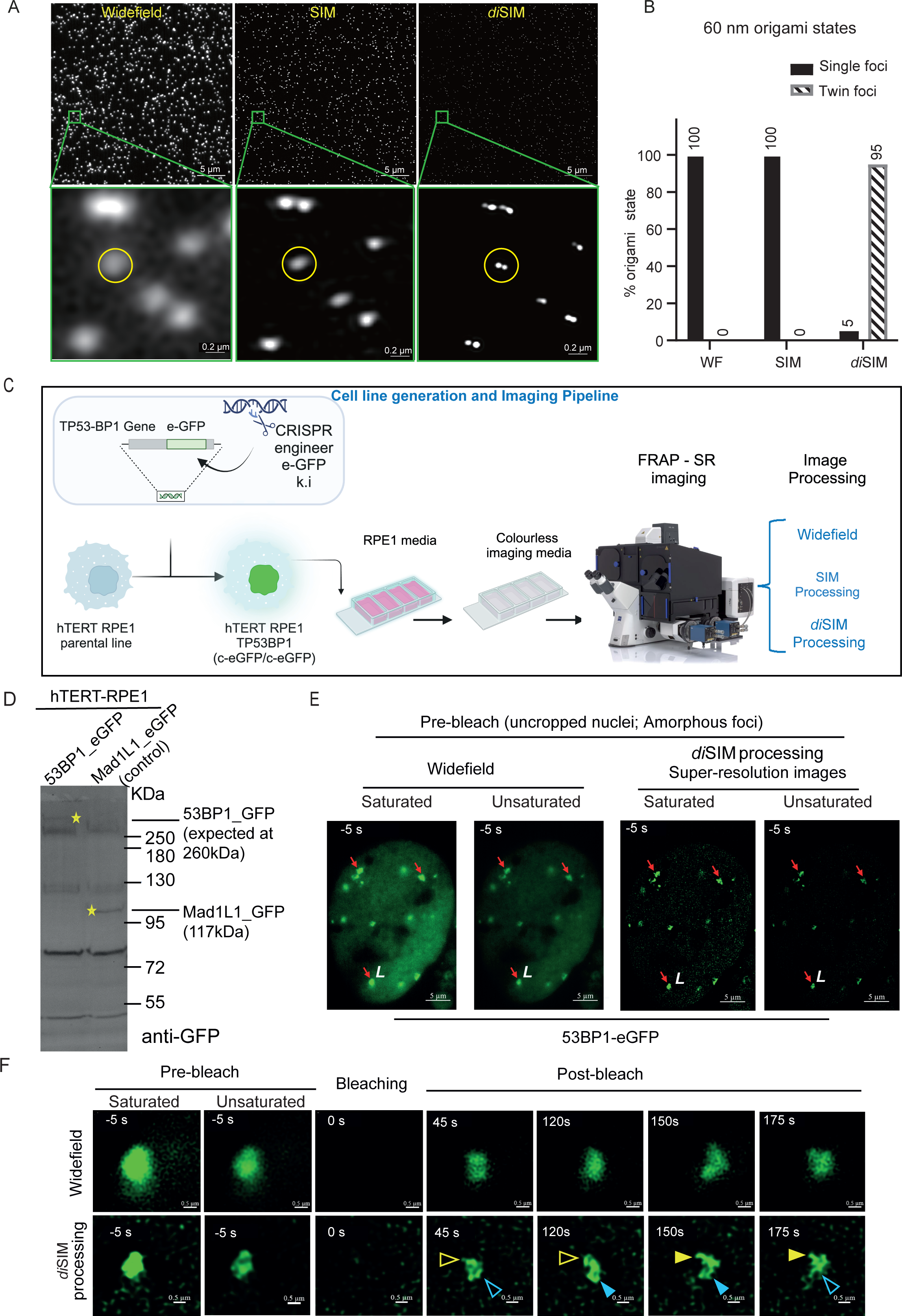
53BP1 foci can be super-resolved into subcompartments using FRAP in the SR regime. **(A)** Uncropped (upper) and magnified (lower) images of 60 nm origami beads imaged and processed for super-resolution using SIM or *di*SIM (*dual iterative* SIM) algorithms as indicated. Scale bars as indicated. Green squares on upper row correspond to magnified crops in the lower row **(B)** Graph shows 60 nm origami bead status either unresolved as a ‘single foci’ or resolved into ‘twin foci’ in unprocessed widefield (WF) images and super-resolved images (using different processing methods) as indicated. **(C)** The illustration shows an experimental design to conduct FRAP of 53BP1-eGFP foci in the SR regime. Using CRISPR/Cas9, an in-frame sequence encoding Green Fluorescent Protein (eGFP) was inserted in the endogenous TP53BP1 loci of the hTERT-RPE1 cell line to introduce a C-terminal eGFP tag. Cells were grown in RPE1 growth medium (DMEM:F12) and moved to a colourless medium (Leibovitz’s L-15 media) for live-cell imaging. SIM and FRAP were conducted using a ZEISS Elyra 7 microscope equipped with a Rapp OptoElectronics photomanipulation module. Images were processed with widefield reconstruction and the sum intensity of Z-stack was used for FRAP kinetic measurements, or SIM and *di*SIM -processed for foci structure analysis and size measurements in the SR regime. **(D)** Immunoblot of lysates of RPE1 53BP1-eGFP or MAD1L1-eGFP (as indicated) probed with anti-eGFP antibodies show the expression of 53BP1-eGFP or Mad1L1-eGFP as expected. RPE1 Mad1L1-eGFP lysate is used as a control. For markers, see a composite pseudo-coloured image presented in Supplementary Figure 1B. **(E)** Images of a RPE1 53BP1-eGFP nucleus showing 53BP1 foci (red arrows) with an amorphous and irregular contour. Widefield and *di*SIM-processed images are shown. The letter ‘L’ in white marks the amorphous foci bleached and tracked for recovery in panel **F**. Scale bar as indicated. **(F)** Cropped time-lapse images of areas marked by a white arrow in **E** show fluorescence intensities before and after bleaching. Recovery images show uneven recovery across the amorphous 53BP1 foci (empty and filled colour arrows show the absence and presence of GFP signal intensities, respectively). Saturation levels were set up for post-bleach recovery images (unsaturated pre-bleach images included). Scale bar as indicated.

53BP1-eGFP foci undergo phase-separation^10,11^ and facilitate long-range interactions between DNA ends^13^. Cellular levels of 53BP1 can influence the protein’s accumulation on DSB sites. To avoid bias from ectopic 53BP1 protein expression, we used CRISPR/Cas9 to fluorescently tag 53BP1 in the hTERT-RPE1 cell line (Figure 1C, Supplementary Table 1). We first characterised the eGFP knock-in cell line with immunoblotting of RPE1 53BP1-eGFP cell lysates using an anti-eGFP antibody, which showed a 260 kDa band corresponding to 53BP1-eGFP - this band was absent in RPE1 Mad1-eGFP cell lysates that displayed a 117kDa band corresponding to Mad1-eGFP as expected (Figure 1D and Supplementary Figure 1B). Next, using Lattice Light Sheet microscopy^24^, we imaged RPE1 53BP1-eGFP cells for 24 hours once every 5 minutes. We confirmed both the steady nuclear levels of 53BP1-eGFP (Supplementary Movie 1) and the typical cell cycle-regulated pattern of 53BP1 foci disappearing in mitosis (except on kinetochores^25^) and reappearing in G1-phase nuclei as G1 bodies^15^ (Supplementary Movie 2; Supplementary Figure 1C), demonstrating that 53BP1-eGFP localises similarly to endogenous 53BP1.

The widefield reconstruction of Lattice SIM movies of RPE1 TP53BP1-eGFP cells showed that some of the 53BP1-eGFP foci presented an amorphous contour that appeared to stretch and collapse loosely (red arrows, Figure 1E, Supplementary Movie 3). Upon SIM processing, these foci which we term ‘amorphous foci’ displayed a dynamic, irregular shape which was not readily evident, although present in Widefield unprocessed images (Compare Supplementary Movies 3, 4a and 4b). To further improve SIM outcomes beyond the 120 nm lateral resolution, we employed computational methods (dual-iterative SIM, *di*SIM, commercial name: SIM^2^) that offered further improvement^22^. In both SIM and *di*SIM processed time-lapse movies, some (but not all) 53BP1-eGFP foci display an amorphous contour and an irregular shape (n=10 cells, Figure 1E and Supplementary Figure 2, Supplementary Movie 4a and 4b for SIM and *di*SIM processing).

Next, we photobleached 53BP1-eGFP foci using a 473 nm laser to analyse the recovery of eGFP fluorescence. For this purpose, time-lapse images were acquired once every 5 seconds for 2 minutes using a leap mode yielding 9 Z-stack. In *di*SIM-processed images, we observed a non-uniform recovery within the foci, leading to changes in positions of eGFP intensity peaks, suggesting different protein diffusion rates within a single 53BP1 foci, indicating distinct subcompartments (Figure 1F). Thus, combining FRAP and SR imaging has the potential to reveal subcompartments of differing protein mobilities within larger subcellular structures.

### Few 53BP1 foci remain compact without separating into amorphous foci

To assess differences in 53BP1 protein mobilities, we conducted fluorescence recovery after photo-bleaching (FRAP) measurements of 53BP1 foci in several cells and calculated FRAP recovery rates using raw unprocessed images. Nuclear foci that appeared as a compact spot in prebleach images were first used for the study (Figure 2A). *di*SIM processed and unprocessed images of compact foci showed uniform recovery as a compact structure during the 2 minutes of FRAP period, and these were analysed for 53BP1-eGFP recovery kinetics (Figure 2B, Supplementary Movies 5 and 6). In these compact 53BP1 foci, 53BP1-eGFP recovery kinetics (mean half time of recovery t_1/2_=19.67 seconds; n=10 cells (Figure 2C)) are similar to what has been reported^10^. In compact foci, 55% of 53BP1-eGFP had recovered within a minute, showing 53BP1 protein exchange (Figure 2C). These studies show that the FRAP rates of 53BP1-eGFP foci in our unprocessed SIM images are comparable to previous non-SIM studies. We conclude that 53BP1, within compact foci, recovers uniformly as a single compartment.

**Figure 2:**
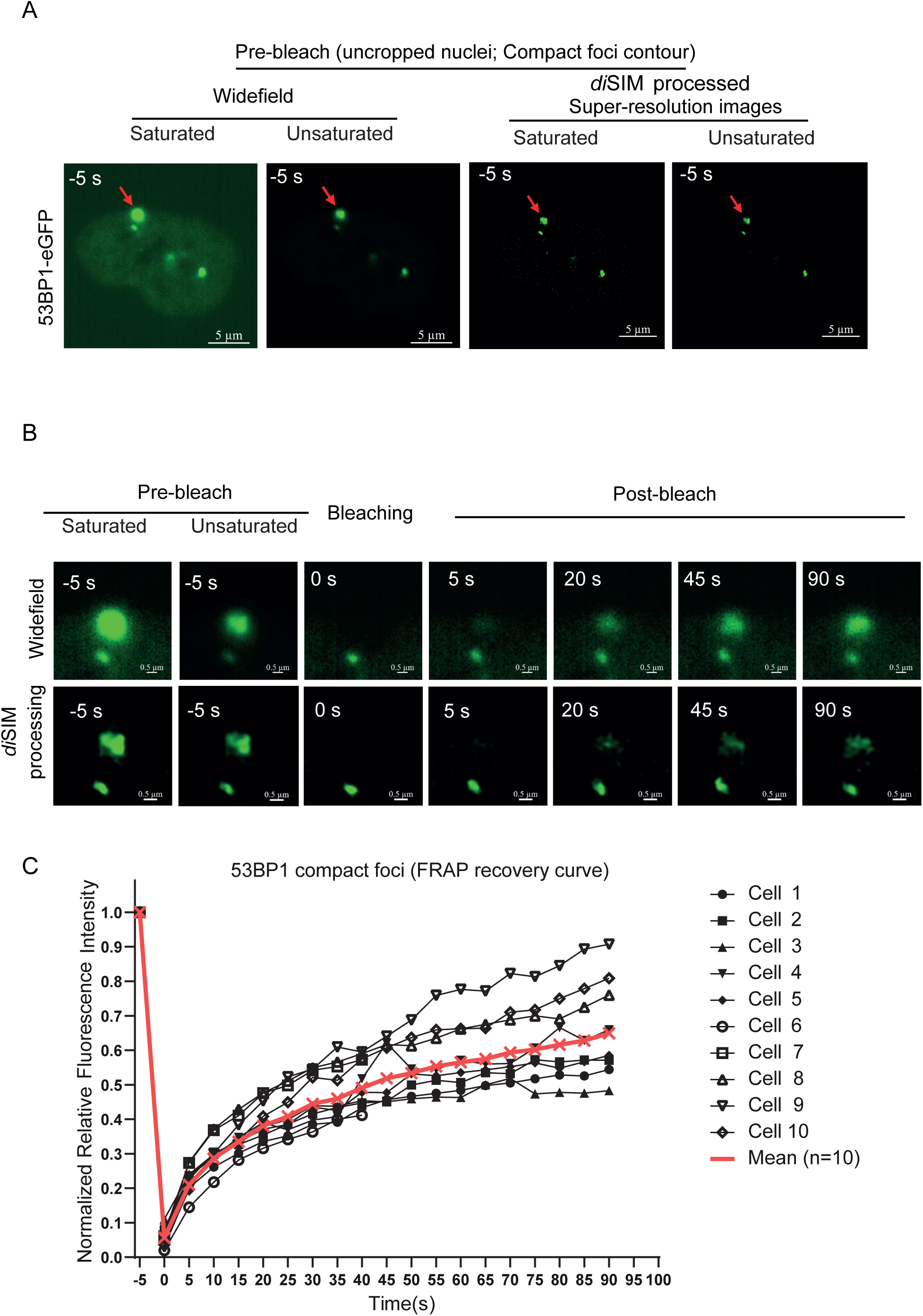
Single compact foci show a uniform recovery of 53BP1-eGFP signal. **(A)** Representative widefield, and *di*SIM-processed prebleach images of nuclei with a compact foci (red arrow). Scale bars as indicated. **(B)** Cropped time-lapse images (widefield or processed *di*SIM as indicated) show 53BP1 foci with a single compact contour photobleached to study FRAP rates. Scale bar as indicated. **(C)** Graph of relative fluorescence intensity of single compartment 53BP1-foci shows FRAP recovery during the 3-minute imaging period following photobleaching. Saturation levels were set up for post-bleach recovery images (unsaturated pre-bleach images included).

### FRAP of 53BP1-eGFP indicates at least two types of 53BP1 foci

We next focussed on 53BP1 foci that present amorphous and dynamically varying contours visible in both unprocessed and SIM-processed movies (Figure 3A, Supplementary movies 7 and 8). We tested whether the FRAP rates of 53BP1 within the amorphous foci are similar to those in the compact foci. Following photobleaching, SIM-processed images showed differential recovery within the 53BP1 foci, with some regions (subcompartments) recovering faster than others (purple arrow, Figure 3B). The mean recovery kinetics of 53BP1-eGFP in the amorphous foci structures showed a half-time of recovery t_1/2_=15.58 seconds; n=10 cells. Occasionally, cells displayed both types of 53BP1 foci (compact and amorphous) within the same nuclei (Supplementary Figure 2). There was no significant increase in 53BP1 foci numbers during imaging (Supplementary Figure 3A and 3B), suggesting no significant DNA damage was introduced during the FRAP-SR imaging session.

**Figure 3:**
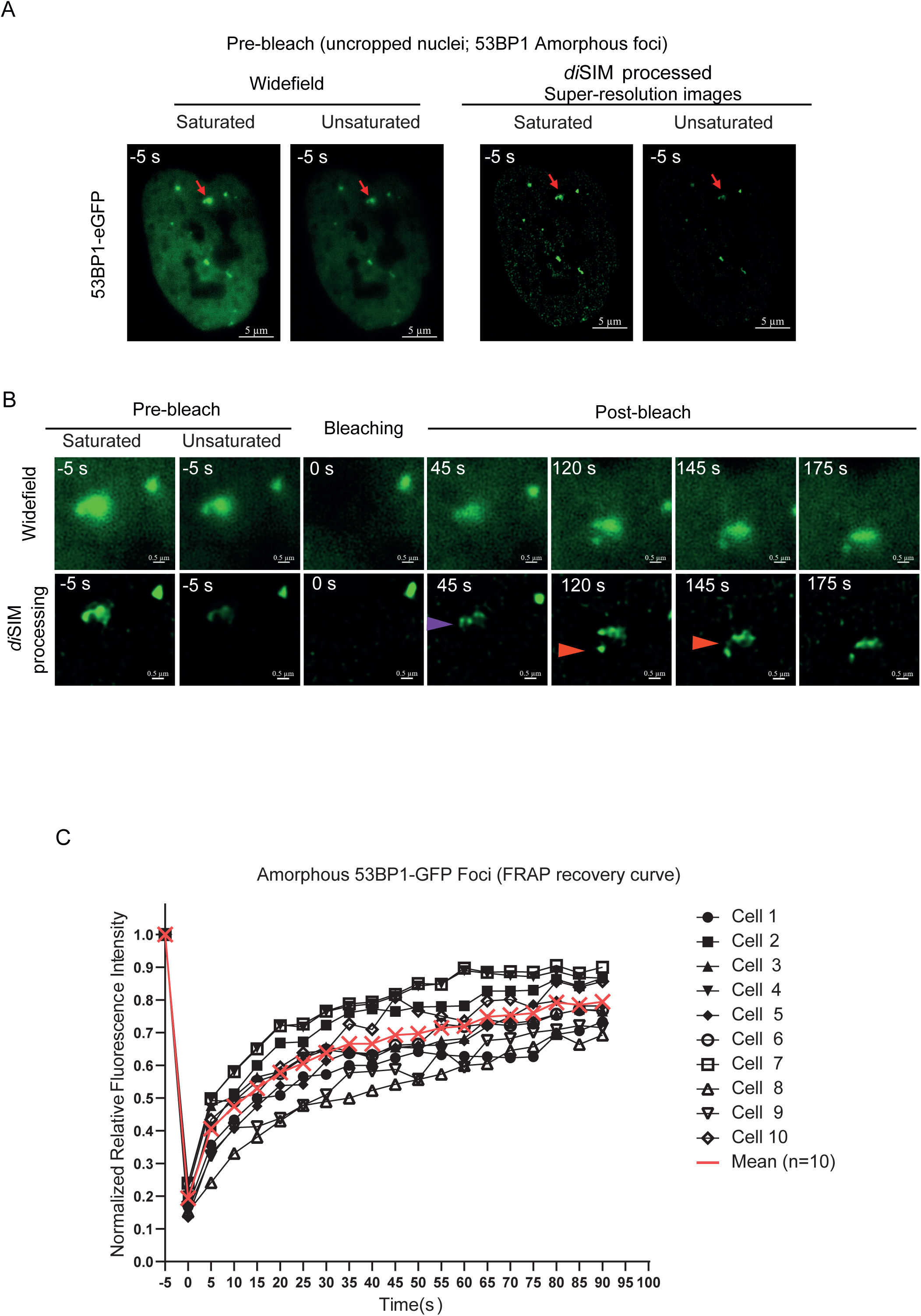
Amorphous 53BP1 foci show subcompartments and nonuniform recovery. **(A)** Representative, unprocessed, SIM or *di*SIM-processed pre-bleach images of nuclei with amorphous foci (red arrow). Scale bar as indicated. **(B)** Cropped time-lapse images (raw or processed (SIM or *di*SIM as indicated)) show a multi-compartment 53BP1-foci that was photobleached to study FRAP rates. Purple arrowhead marks compartments that recover earlier, and red arrowhead tracks foci morphology and position change. Scale bar as indicated. **(C)** Graph of relative fluorescence intensity of multi-compartment 53BP1-foci shows FRAP recovery during the 3-minute imaging period following photobleaching. Saturation levels were set up for post-bleach recovery images (unsaturated pre-bleach images included).

In summary, we conclude that 53BP1 foci can be separated into at least two different types, based on a) the amorphous or compact foci contour and b) the presence of subcompartments with differing 53BP1 protein mobilities.

### 53BP1 foci displaying subcompartments may be more active than others

FRAP rates of 53BP1-eGFP are more heterogeneous in compact foci than amorphous foci, suggesting varying protein mobilities^26^, although there was no statistical significance between the two groups (Figure 4A). We set out to test if any other subcellular differences can be correlated to explain the subcompartments we observe in amorphous foci, but not in compact 53BP1 foci.

**Figure 4:**
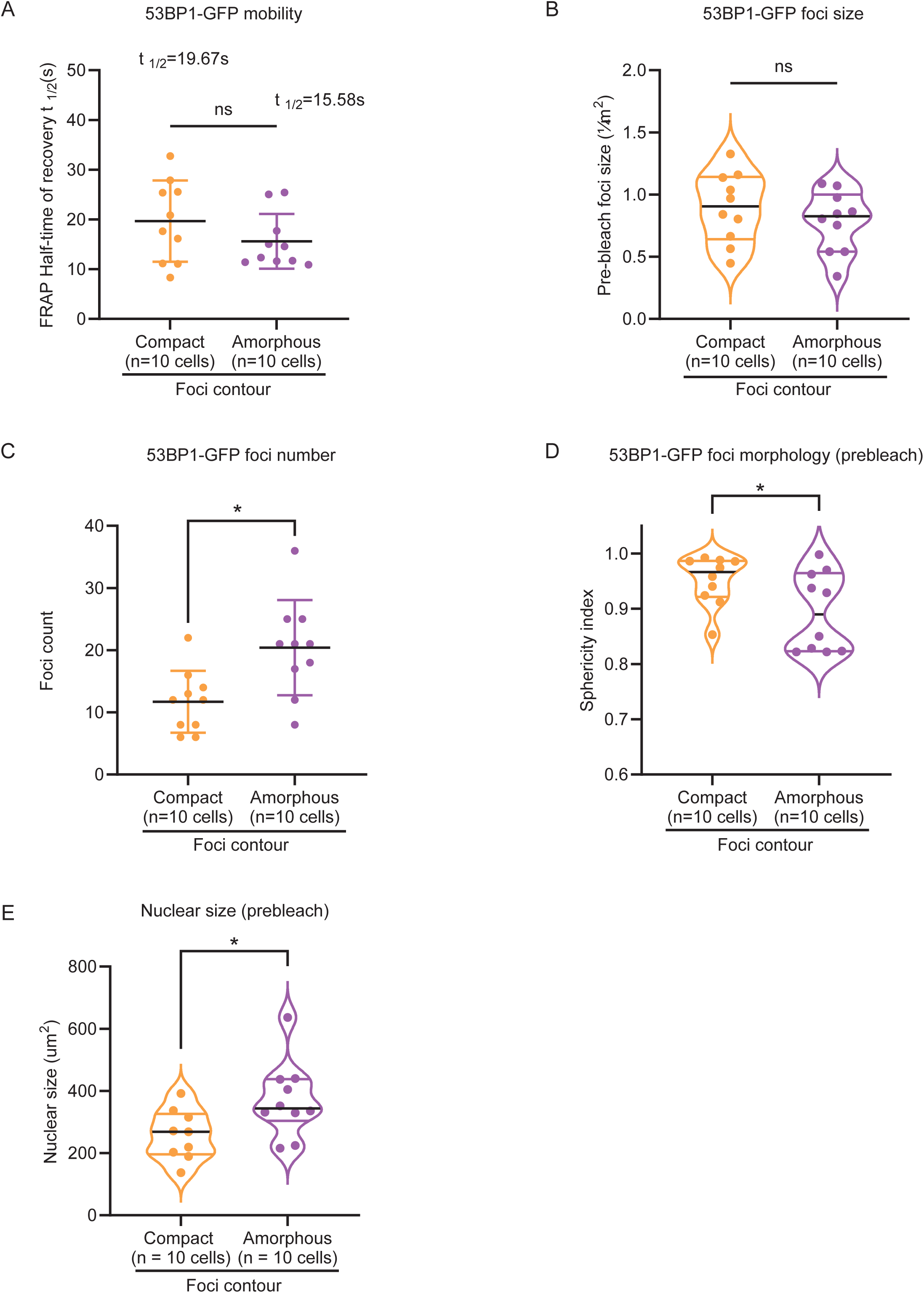
53BP1 foci contours, compact and amorphous, show an association with nuclear but not foci sizes. **(A)** Graph showing the distribution of half-times of 53BP1-eGFP FRAP recovery indicates an increased heterogeneity in recovery rates of single compact foci compared to amorphous foci. Two-tailed Mann-Whitney test, p=0.3527, no significant difference). **(B)** Graph showing no significant difference in foci size (measured as area) between compact foci and amorphous foci (pre-bleach) in *di*SIM-processed images of RPE1 53BP1-eGFP cells (Two-tailed Mann-Whitney test, p=0.4359, no significant difference). **(C)** Graph showing some difference in 53BP1 foci numbers (measured as counts in pre-bleach nuclei) of SIM-processed images of cells displaying either compact foci or amorphous foci (Two-tailed Mann-Whitney test, p=0.1803, no significant difference). **(D)** Graph showing the sphericity index of 53BP1 foci (morphology of prebleach foci was considered for this analysis (Unpaired t-test, p=0.0459, no significant difference). **(E)** Graph showing the distribution of nuclear sizes in cells corresponding to 53BP1 foci studied for FRAP rates in **A (**Two-tailed Mann-Whitney test, p=0.0279, significant difference; 1 cell with an unusually large nucleus was removed from the compact group). Median values are marked using black lines and quartiles are marked using colour lines.

OPT domains, 1.3 microns in diameter, appear in the G1 phase and are thought to disappear in the S phase^15,27^, so first we analysed the sizes of the foci we photobleached. Both compact and amorphous foci exhibit a wide range of sizes from 0.34 to 1.33 micron^2^ as measured using *di*SIM-processed images. We find no significant difference in the sizes of the compact and amorphous foci (Figure 4B), showing that their activities may not strictly depend on the size of phase-separated structures.

53BP1-associated chromatin and DSB mobility has been reported ^28–30^. To test the extent of mobility in compact and amorphous foci, we measured foci displacement using centroids of the 53BP1 foci before and after photobleaching (to assess movement within 40 seconds) (Supplementary Figure 4A). Five of the ten compact foci showed no displacement (Supplementary Figure 4B and 4C, n = 20 cells). Although we did not find any statistical difference between the displacement of amorphous and compact foci, 50% of compact foci did not display any mobility, distinguishing the mobility likelihood of compact and amorphous foci.

Soon after mitosis, very few 53BP1 foci (as G1 bodies) are expected to arise from the previous cell cycle, whereas in the S-phase many more 53BP1 foci are expected^6,14,31^. Therefore, we explored whether the number of foci within the nucleus differs in the cells that display compact or amorphous foci. Segmentation of images to automatically count 53BP1 foci based on eGFP intensities showed a moderate reduction in the total number of 53BP1 foci in cells with compact foci compared to those with amorphous foci (n=20 cells) (Figure 4C). Next, we plotted the sphericity index to characterise the spikiness property of the foci’s exterior boundary. We found that compact foci tended to be more spherical compared to amorphous foci (Figure 4D). Lastly, we correlated the size of nuclei associated with amorphous or compact foci in an unbiased manner showed a median 1.1-fold reduction in the nuclear size of cells displaying compact foci compared to amorphous foci (Figure 4E), suggesting cell cycle-associated changes. Together, these quantitative studies show clear differences between 53BP1 foci contour, displacement propensity and numbers but not foci sizes, suggesting a closer link between nuclear size and foci contours compared to foci size *per se*.

### Lattice light-sheet movies show amorphous foci amidst multiple 53BP1 foci induced following aphidicolin treatment

Nuclear size changes through the cell cycle. To test whether differences in 53BP1 foci morphology are associated with the cell cycle, we characterised 53BP1 foci occurrence and resolution times in long-term live-cell movies of aphidicolin-treated and -released cells. Using a lattice light-sheet microscope that offers a gentle illumination profile, we imaged RPE1 53BP1-eGFP once every 5 minutes for up to 24 hours. Following aphidicolin treatment, we observed three types of nuclei based on 53BP1 foci pattern: (i) no prominent foci (termed ‘no foci’), (ii) one or two large compact foci or (iii) several diffused and small foci (termed multi-foci) (Figure 5A). 360 minutes after aphidicolin release, 30% of nuclei displayed ‘no foci’, 58% of nuclei displayed ‘multi-foci’ and 12% of nuclei displayed ‘compact foci’ (Figure 5B).

**Figure 5:**
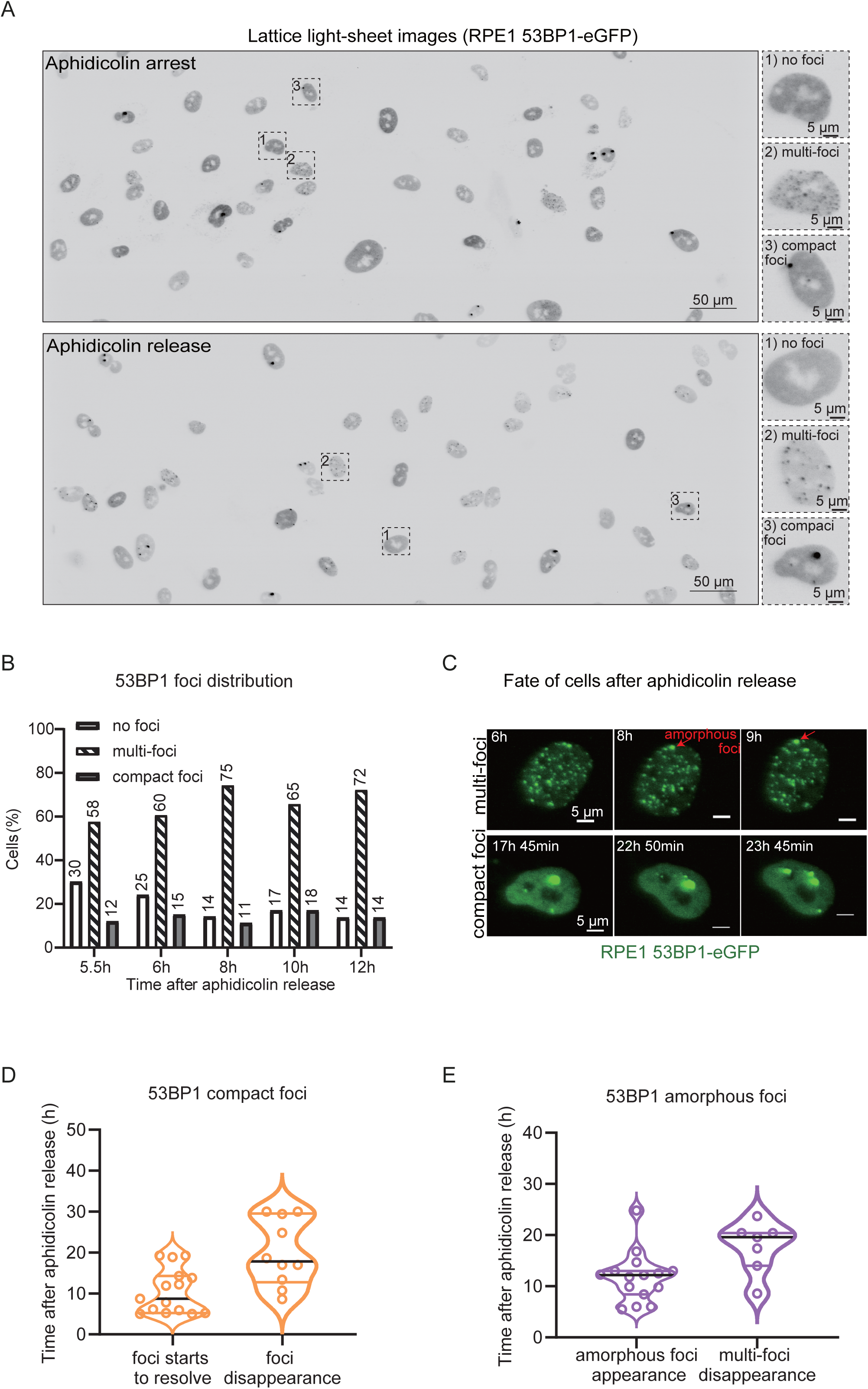
Lattice light-sheet (LLS) imaging shows faster resolution of amorphous versus compact 53BP1 foci. **A)** Lattice light-sheet images of cells released from aphidicolin treatment or retained in aphidicolin during imaging. Nuclei displaying no 53BP1 foci, compact foci or amorphous foci are shown in magnified crops. Scale bar as indicated. **B)** Bar graph showing the percentage of cells displaying no 53BP1 foci (no foci), multiple foci throughout the nuclei (multi-foci) or few compact foci. Numerical values on top of bars indicate percentage values (n = 33 to 36 cells / time frame). **C)** Images show changes in the fate of foci in nuclei displaying multi-foci or compact foci. Amorphous foci showing dynamic changes in shape are marked in red. **D)** Violin plot showing the beginning and end of compact foci resolution times following a release from aphidicolin treatment in cells expressing 53BP1-eGFP imaged using lattice light-sheet microscopy as in **A. E)** Violin plot showing the appearance of amorphous foci and disappearance of multi-foci state after aphidicolin treatment release in movies of RPE1 cells expressing 53BP1-eGFP as in **A.** Median values are marked using black lines and quartiles are marked using coloured lines.

Dynamic changes in the number and size of compact foci were observed over time (Figure 5C). Amorphous foci were observed in nuclei displaying a ‘multi-foci’ pattern (Figure 5C). Quantifying compact foci resolution times showed that the foci started to resolve as early as 5 hours after aphidicolin release but showed a wide range of foci resolution times (Figure 5D). Similarly, G1-bodies that occur soon after mitosis and appear compact showed a range of foci resolution times (Supplementary Figure 5). In multi-foci nuclei, we observed the appearance of amorphous 53BP1 foci but could not track its disappearance due to resolution limitations in lattice light-sheet microscopy (Figure 5E). However, we find that the multi-foci state of nuclei disappeared completely with time, suggesting cell cycle regulation (Figure 5E). In summary, the timing of foci appearance and foci resolution in cells released from aphidicolin suggest that 53BP1 foci morphology may reflect different protein activities or functional states of the foci through the cell cycle.

### FRAP-SR of amorphous foci in aphidicolin-released cells display subcompartments of rapid 53BP1 recovery

Using FRAP-SR, we compared 53BP1-eGFP recovery in compact and amorphous foci of aphidicolin-arrested or -released cells (Supplementary Figure 6A). In aphidicolin-arrested cells, we measured 53BP1 recovery only in compact foci (amorphous foci were not studied due to the crowding of multiple foci). In aphidicolin release conditions (4-8 hours after release), we measured 53BP1 recovery in compact and amorphous foci (Supplementary Figure 6B). In the aphidicolin-release state, 53BP1-eGFP in amorphous foci recovered with a mean half-life of 10.25 s (Figure 6B). In aphidicolin -arrested or -release states, compact foci displayed a 53BP1 half-life of nearly 28.6 s (Figure 6B), suggesting a slower 53BP1 protein exchange than compact foci in untreated cells (Figure 4A). Faster 53BP1 recovery rates in amorphous compared to compact foci following aphidicolin release (Figure 6B) indicate accelerated 53BP1 exchange in the subcompartments. These were also reflected in immobile factions across conditions (Supplementary Figure 6C).

**Figure 6:**
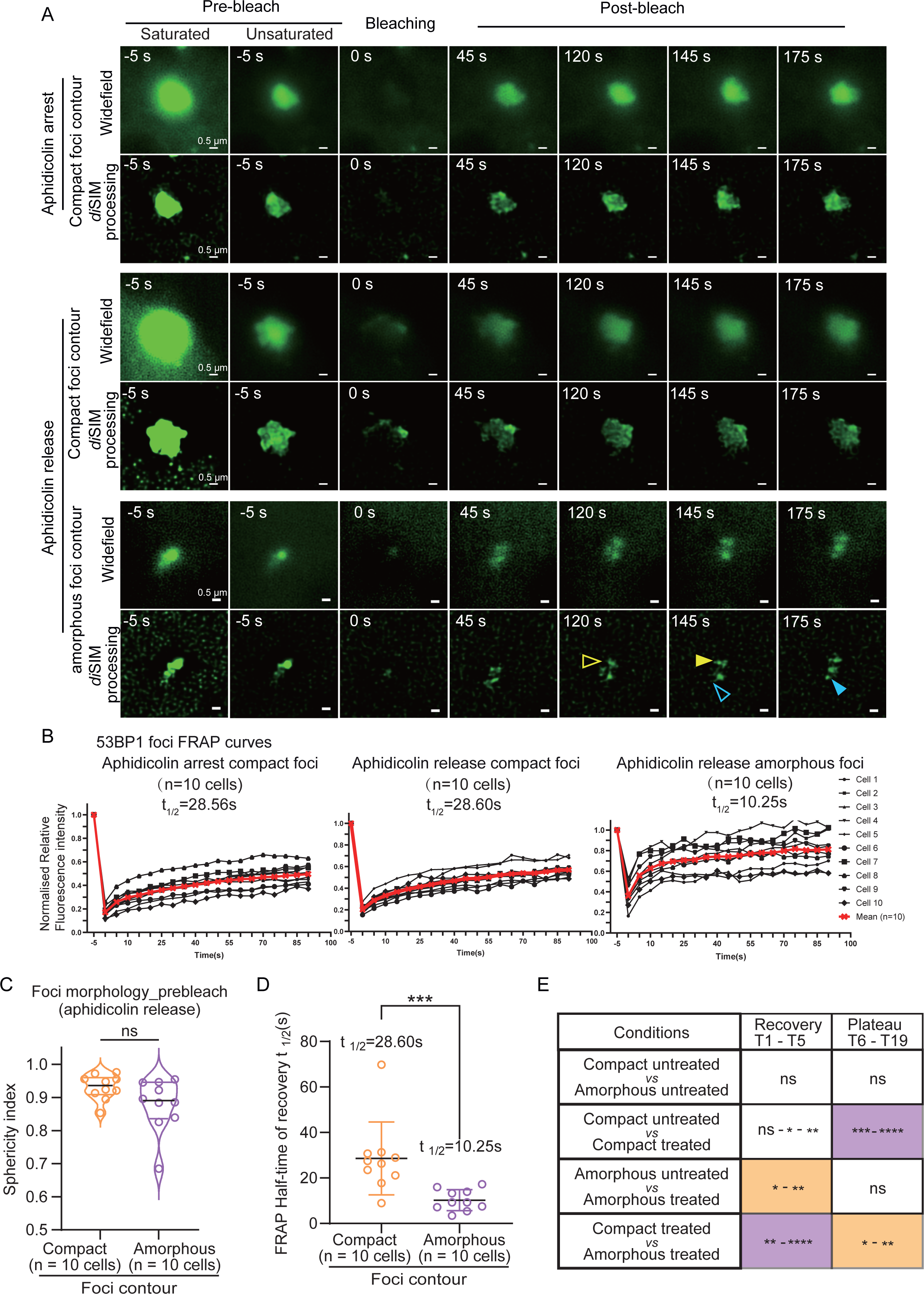
Rapid 53BP1 protein recovery in amorphous compared to compact foci following aphidicolin treatment. **A)** Cropped time-lapse images of compact or amorphous foci as indicated in aphidicolin-arrested or released cells. Saturation levels were set up for post-bleach recovery images (unsaturated pre-bleach images included). Post-recovery images (120 - 175 s) of amorphous 53BP1 foci continue to show uneven eGFP recovery (empty and filled colour arrows mark the absence and presence of signal intensities, respectively). Scale bar as indicated. **B)** Graphs of relative fluorescence intensity of 53BP1-foci show FRAP recovery during the 3-minute imaging period following photobleaching in compact or amorphous foci in aphidicolin-arrested or -released cells as indicated. t_1/2_ indicates half maximal recovery times for each condition. **C)** Violin plot showing sphericity indices of compact and amorphous 53BP1 foci, under prebleach conditions, in RPE1 cells released from aphidicolin treatment (Two-tailed Mann-Whitney test, p=0.0524, no significant difference). **D)** Graph showing the distribution of half-times of 53BP1-eGFP FRAP recovery indicating a faster recovery rate in amorphous foci compared to compact foci (Two-tailed Mann-Whitney test, p=0.0003, significant difference). E) Table shows statistical differences extracted by comparing FRAP recovery curves in two conditions indicated during recovery or plateau period as indicated in Supplementary Figure 6C. Two way ANOVA was used to identify nonsignificant (ns) or significant differences among groups (***** P≤ 0.00001, **** P≤ 0.0001,*** P≤ 0.001, * P≤ 0.05 and ns P> 0.05). Median values are marked using black lines and quartiles are marked using colour lines.

Although normalisation of FRAP curves to assess immobile fraction was difficult due to varying intensities of the foci, comparing normalised FRAP mean values across 10 cells showed that the immobile fraction in aphidicolin-arrested compact foci is the largest compared to untreated amorphous foci which displayed the smallest immobile fraction across the tested conditions (Supplementary Figure 6C). We observed no significant differences between the foci morphology, measured as sphericity index (Figure 6C). However, the FRAP half-time recovery rates are significantly different between compact and amorphous foci in aphidicolin release conditions (Figure 6D). We next compared the differences in FRAP rates of 53BP1-eGFP foci during the eGFP signal recovery period (T1 to T6 time frames, less than 25 s after photobleaching) and the signal plateau period (T7-T19 time frames, more than 25 s after photobleaching) (Supplementary figure 6C). In alignment with half-time measurements (Figure 6C and 4A), compact and amorphous foci in aphidicolin treated but not untreated cells responded significantly differently during the signal recovery period (Figure 6E). In contrast, compact foci in aphidicolin treated *versus* untreated cells showed a significant difference in the recovery period (Figure 6E). These findings indicate differences in 53BP1 exchange, foci contours and subcompartments in aphidicolin treated and untreated cells which may reflect differences in 53BP1 protein activities.

Combining FRAP and SR with additional DNA damage repair proteins would allow a deeper exploration of how DDR machinery access and occupancy are regulated. Collectively, the quantifications show the strength of FRAP-SR microscopy to compare differences in 53BP1 foci contours, protein diffusion rates within subcompartments, and foci mobility for rigorously exploring protein activities and function within subcellular structures.

## Discussion

We combine FRAP and SR microscopy studies to investigate 53BP1 protein exchange rates and subcellular structural changes in the SR regime, taking a step beyond SR or FRAP studies done separately. Our SR live-cell studies of 53BP1 foci in unperturbed cells indicate that the foci can either remain compact or amorphous. Amorphous foci show dynamic irregular shapes and increased foci movement, suggesting activity. FRAP studies demonstrate that protein mobilities within the amorphous foci can be uneven, leading to partial recovery of 53BP1 foci compartments and revealing subcompartments of differential protein activity (Figure 7). In contrast, 53BP1-eGFP in compact foci recovers uniformly. Following aphidicolin treatment, 53BP1-eGFP recovers faster in amorphous than compact foci, indicating differences in 53BP1 protein activities. In addition to showcasing the strengths of FRAP-SR, our findings have conceptual implications on whether other components of the DDR machinery are uniformly found within the 53BP1 foci and whether different 53BP1 mobilities indicate differential roles in distinct subcellular events (eg., replication or repair).

**Figure 7:**
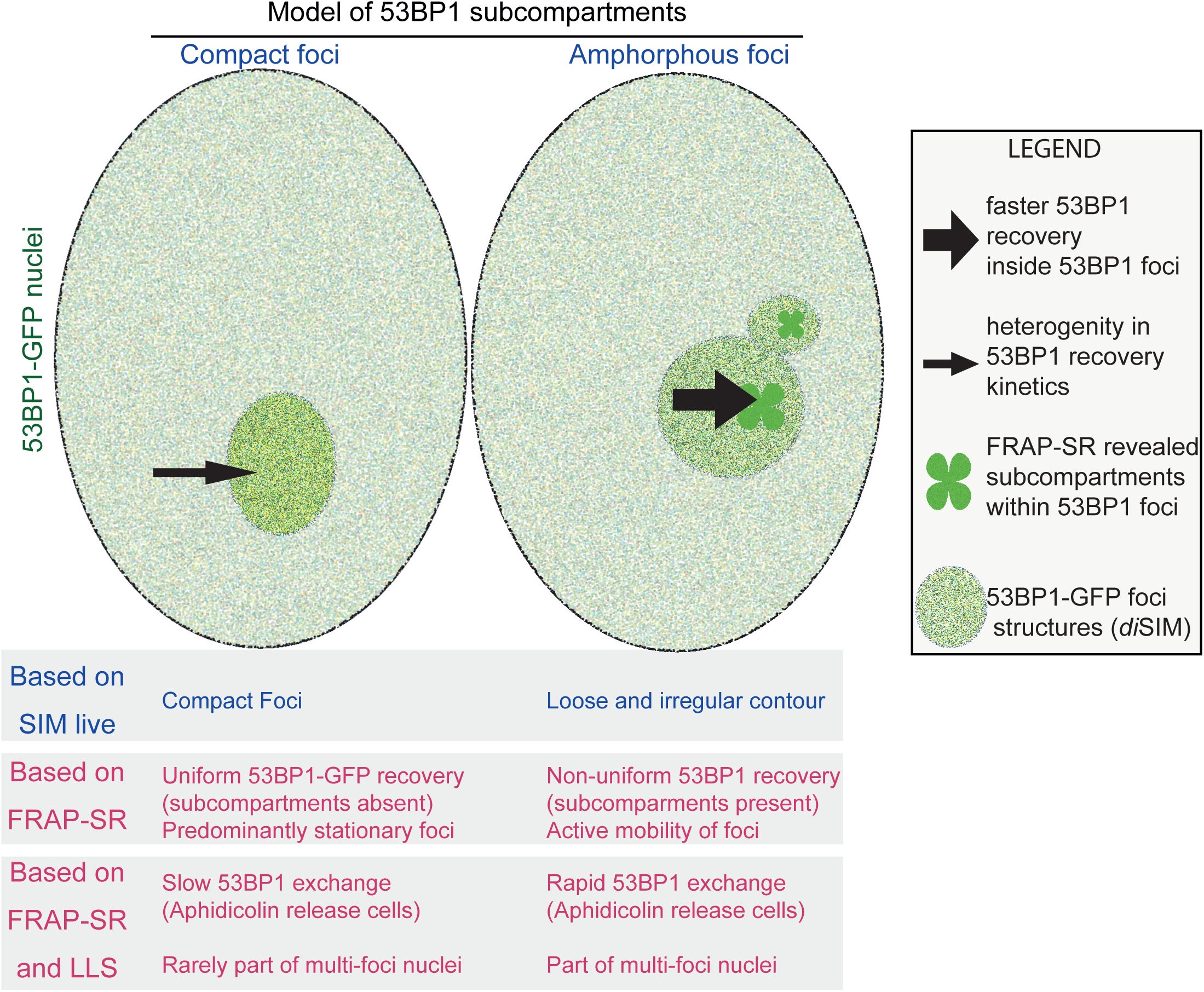
Model of subcompartments within 53BP1 foci showing sites of distinct activities based on differing 53BP1 protein diffusion rates. FRAP in the SR regime shows (i) a slower 53BP1 exchange rate between foci and nucleoplasm compared to exchange within the foci and subcompartments and (ii) heterogeneity in the recovery of eGFP-53BP1 signals following photobleaching of compact compared to amorphous foci. Differing protein mobilities and super-resolved structural dynamics may reflect distinct roles of 53BP1 activities and functions as they correlate with different foci resolution kinetics. FRAP-SR can extend the imaging toolset to probe the dynamic regulation of subcellular structures.

53BP1 foci can arise from different types of replication stress, clustered DNA damage or non-DNA damage sites. SMLM studies have shown the existence of subfoci of 53BP1 following DNA damage, which arises from clustered DNA damage^32^. Although 53BP1 has been shown to undergo phase separation^10,11^, there was no prior knowledge of distinct protein mobilities within 53BP1 subcompartments which we report here. Long-term imaging of cells after a FRAP-SR cycle could help measure and model foci and protein mobilities to explore whether compact foci follow sub-diffusive dynamics. Our FRAP-SR studies suggest that within phase-separated 53BP1 foci, there can be activity centres with increased protein mobility, which appear as subcompartments. Consistent with this model, in cells released from aphidicolin treatment, amorphous foci show faster 53BP1 protein exchange (FRAP-SR) compared to compact foci (Figure 6).

FRAP-SR studies using SIM are strategically placed to resolve and link dynamic protein interactions and subcellular structural changes within a range of 60 - 120 nm without causing phototoxicity. High-throughput microscopy studies comparing the localisation of isoforms, variants and mutants of proteins across the cell cycle can benefit from a single-step framework to explore protein mobilities/interactions in addition to changes in the organelle they decorate^33^. Structural changes due to frequent fusion and rare fission events have been reported to explain the droplet-like behaviour of 53BP1 foci^10^ wherein nuclear bodies and IR-induced foci recover with similar kinetics, suggesting similar diffusion rates of 53BP1 molecules. In study of aphidicolin treated cells from different cell cycle stages, we find differences in the recovery kinetics of 53BP1 within compact and amorphous foci, indicative of distinct protein interactions and associated functions in these compartments. Consistent with these findings, functional interrogation of DNA damage response variants showed 53BP1 mutants to modulate protein binding without affecting damage response^34^.

DNA break repair studies have shown a fast component for resolving most of the breaks and a slow component for resolving some of the breaks^6,35–37^. Whether the two different types of super-resolved 53BP1 foci indicate fast and slow resolving breaks (differing 53BP1 activities) can be confirmed with FRAP-SR of other DDR-associated proteins. Alternatively, a dynamic amorphous foci could indicate spaces of higher mobilities within the foci, consistent with BRCA1-mediated active exclusion of 53BP1 from DNA repair sites proposed in previous SIM studies of BRCA1 and 53BP1^17,19^. In our studies using unperturbed RPE1 cells, we ruled out foci arising from incomplete DNA replication or replication stress; these can be triggered by aphidicolin, which increases the incidence of 53BP1 foci in the G1 phase of the next cell cycle^15^.

High-speed nanoscale imaging of live cells has provided new insights (eg., organelle contact sites^38^ or induction of DSBs following radiation^29,39^). Also, high-speed volumetric studies of live cells enable the probing of subcellular movements across mesoscales^40^. Using Lattice SIM here, we combine volumetric SR methods and FRAP to reveal that not all 53BP1 foci show similar protein mobilities and that foci that dynamically resolve into multiple compartments show increased 53BP1 protein mobility (Figure 7). Based on aphidicolin treated and untreated cell outcomes, we propose that differing 53BP1 protein mobilities and foci structures may conceptually relate to different roles of 53BP1 within the foci, a mechanism well suited for varied spatiotemporal events. 53BP1 is involved in both rapid and slow cellular events: resolving DNA double-strand breaks^39,41^ and fast chromatin mobilities^29^ are in the order of seconds and minutes while 53BP1’s role in mitosis and replication stress^42^ or heterochromatin binding^11^ are in the order of minutes to hours requiring different types of interactors and regulatory mechanisms. Our foci displacement studies show that 75% of 53BP1 foci are mobile, allowing the foci to support long-range interactions. Long-range joining of DNA breaks, as in the distal joining of V-DJ mediated by 53BP1, are important as defective cells can experience extensive degradation of the unrepaired coding ends^13^, leading to genomic instability. In the long-term, combining FRAP and *di*SIM can take us closer to exceeding current limitations in studying photosensitive subcellular structures with varied protein mobilities, activities and roles.

## Supplementary Information

### Supplementary Figures

**Supplementary Figure 1: Characterisation of *di*SIM lateral resolution and CRISPR-engineered hTERT-RPE1 53BP1-eGFP cell line.**

**A)** Distribution of distance between the twin foci observed following *di*SIM processing of 60 nm origami beads (data relates to images in Figure 1A). **B)** Immunoblot of lysates of RPE1 cells CRISPR engineered either at the TP53BP1 or MAD1L1 gene locus to introduce a C-terminal eGFP tag. Immunoblots probed with anti-eGFP antibodies (pseudo-coloured in green) show the expression of 53BP1-eGFP or Mad1L1-eGFP as expected. In red are protein marker lanes with estimated molecular weights marked on the right. Mad1L1-eGFP lysate is used as a control. The immunoblot is related to the grayscale image presented in Figure 1B. **C)** Time-lapse LLS7 images show 53BP1-eGFP foci appearing normally as G1 bodies soon after mitosis. Images of a G2 phase interphase cell becoming mitotic and disassembling 53BP1 foci (Figure related to Supplementary Movie 1). Yellow arrowheads mark G1 bodies.

**Supplementary Figure 2: Compact and Amorphous foci 53BP1-eGFP foci can occasionally co-occur in cells**

Super-resolution *di*SIM time-lapse images of an RPE1 53BP1-eGFP interphase nuclei show two types of 53BP1-eGFP foci: right foci remains as compact foci (blue asterisk) while on the left amorphous foci (yellow asterisk) shows dynamic irregularities of foci contour growing and shrinking (red arrows mark compact or amorphous foci). No photobleaching was conducted. Scale bars as shown.

**Supplementary Figure 3: No increase in foci count during FRAP-SR studies**

**A)** Super-resolution *di*SIM images of compact and amorphous foci. Five representative images from the FRAP study (pre- and post-bleaching) were presented to showcase a) full recovery of 53BP1 b) uniform recovery on the left (compact foci) c) multi-compartment foci on the right (amorphous). Scale bars as shown. **B)** 53BP1 foci count, in compact or amorphous foci-bearing nuclei, measured using SIM-processed 53BP1-eGFP images showing no significant increase in foci count following bleaching. Pre-bleach and Post-bleach (5 or 80s after bleaching) foci count shown. Nonsignificant ‘ns’ differences were estimated using two way ANOVA.

**Supplementary Figure 4: Compact foci are frequently stationary compared to amorphous foci**

**A)** *di*SIM processed super-resolution images of compact and amorphous foci. Representative images from the FRAP study show prebleaching, bleached and recovery images. Red arrows mark the bleached foci. **B) and C)** Displacement of centroids of 53BP1 foci between prebleached and recovered status (40 seconds post-bleaching) showed a small but significant increase in the amorphous foci compared to compact foci (n=10 cells, each condition). Pink arrow shows five foci with no discernible displacement. Displacement of photobleached foci in other cells indicated on the right (as legend).

**Supplementary Figure 5: G1 bodies show variable 53BP1-eGFP resolution period. A)** Cropped lattice light-sheet (LLS) microscopy images show G1 bodies that form soon after mitosis in two daughter cells. RPE1 53BP1-eGFP cells were treated with aphidicolin overnight for 10-16 hours and released before 24 hours of imaging. Red arrows mark the G1 body with a long resolution time (>10 h), and red asterisks marks G1 bodies with moderate resolution time (3-7 h). Data representative of 4 independent repeats (n = 34 cells). Scale bar as indicated. **B)** Cumulative frequency (%) graph showing the time taken to resolve G1 bodies in time-lapse movies of cells treated as in **A**. 3 of 34 cells did not resolve foci during the period of imaging. **C)** Violin plot showing median time taken for the resolution of G1 bodies in cells released from aphidicolin treated as in **A**. 3 cells that failed to resolve are omitted from this data. Median value is marked using a black line.

**Supplementary Figure 6: Aphidicolin synchronisation shows foci dynamics across the cell cycle. A)** Aphidicolin treatment and release regime for live-cell imaging. RPE1 53BP1-eGFP cells treated with aphidicolin overnight for 10-16 hours were released and imaged for compact or amorphous foci as indicated. **B)** Uncropped saturated and unsaturated widefield super-resolved images of cells treated as in A. Prebleach images are presented. Scale bar as indicated. **C)** Graph of normalised mean values of FRAP recovery curves to highlight differences in immobile fraction of 53BP1-eGFP in compact or amorphous foci-bearing cells. T6 (25 s) was used as a threshold based on t_1/2_ in Figure 6B, 2C and 3C for recovery and plateau period studies in Figure 6E.

### Supplementary Tables

**Supplementary Table 1: Genomic sequence analysis of RPE1 53BP1-eGFP** Table of PCR sequencing results shows modifications on each allele for clone 213 hTERT-RPE1 - TP53BP1 (C-eGFP/C-eGFP) following sequence analysis of the targeted and non-targeted allele-specific PCR products.

### Supplementary Movies

**Supplementary Movie 1: Long-term lattice light sheet imaging shows steady levels of 53BP1 foci in interphase and mitosis.** Long-term time-lapse images taken once every 5 minutes of a 4-well glass-bottom ibidi^TM^ dish from a single position show 53BP1-eGFP foci (scenes) of RPE1 53BP1-eGFP cell line showing steady levels of 53BP1-eGFP foci in interphase nuclei.

**Supplementary Movie 2: Lattice light-sheet imaging shows cell cycle-associated appearance and disappearance of 53BP1 foci in interphase and mitosis, respectively.** Time-lapse images acquired once every 10 minutes from two distinct positions (scenes) of RPE1 53BP1-eGFP cells show 53BP1-eGFP nuclear signals that disappear during mitosis and reappear in interphase daughter cells. White arrow marks mitotic cell, and orange arrow marks mitosis-associated foci appearance.

**Supplementary Movie 3: Time-lapse movie of unprocessed images of RPE1-TP53BP1-eGFP cells.** Time-lapse images acquired once every 5 sec show a single 53BP1 foci (marked with a red arrow) bleached at 5 secs. Bleached foci recovery fluorescence signal across the foci. The movie corresponds to the cell shown in Figure 1. The foci are resolved into separable compartments following *di*SIM processing as in Supplementary Movies 3a and 3b. Scale bar as shown.

**Supplementary Movie 4a and 4b: SIM and *di*SIM processed time-lapse movie of RPE1-TP53BP1-eGFP cells.** A single 53BP1 foci (marked with red arrow) bleached at 5 sec shows uneven fluorescence recovery across the foci, revealing multiple compartments. Some but not all foci resolve into multiple compartments compared to the raw unprocessed movie (Supplementary Movie 3). Images that underwent SIM processing (4a) and *di*SIM processing (4b) correspond to the images in Figures 1E and 1F. Scale bar as shown.

**Supplementary Movie 5: *di*SIM processed time-lapse movie of a compact compartment 53BP1 foci in RPE1-TP53BP1-eGFP cells.** A single 53BP1 foci (marked with red arrow) bleached at 5 sec shows even fluorescence recovery across the foci, revealing a single compartment. Movie corresponds to the cell shown in Figure 2A and 2B. Scale bar as shown.

**Supplementary Movie 6: Unprocessed time-lapse movie of a compact 53BP1 foci in RPE1-TP53BP1-eGFP cells.** A single 53BP1 foci (marked with red arrow)bleached at 5 sec shows even fluorescence recovery across the foci, revealing a single compartment. The movie corresponds to the cell shown in Figures 2A and 2B. Scale bar as shown.

**Supplementary Movie 7: FRAP-SR of an amorphous 53BP1 foci showing multiple compartments upon 53BP1-eGFP recovery. *di*SIM-processed** images of 53BP1 foci (marked with red arrow) bleached at 5 sec show uneven fluorescence recovery across the foci revealing multiple separable compartments. The movie corresponds to the cell shown in Figures 3A and 3B. Scale bar as shown.

**Supplementary Movie 8: Unprocessed time-lapse movie of an amorphous 53BP1 foci in RPE1-TP53BP1-eGFP cells.** 53BP1 foci (marked with red arrow) bleached at 5-sec show fluorescence recovery across the foci, revealing multiple compartments. The movie corresponds to the cell shown in Figures 3A and 3B. Scale bar as shown.

## Methods

**Cell culture and Media for CRISPR-Engineered RPE1 TP53BP1-eGFP clones** CRISPR/Cas9 was used to engineer the endogenous locus of either TP53BP1 or Mad1_L1 gene of hTERT-RPE1 cell line to integrate an in-frame sequence encoding eGFP. RPE1 53BP1-eGFP (clone 213) cells and RPE1 Mad1_L1 (clone 73) cells were cultured in plastic dishes (Corning 430641U) and grown in DMEM/F12 media (21331-046). For aphidicolin treatment, the cells were treated with 1 microM aphidicolin overnight (10-16 hours). The aphidicolin release group was washed 5 times with 15-20 minutes of incubation at 37 Celsius and then transferred to Leibovitz’s L15 medium for imaging. The aphidicolin arrest group was transferred to Leibovitz’s L15 medium with 1 microM aphidicolin.

**Lattice Lightsheet (LLS7) Imaging** For live-cell imaging using LLS7 (Lattice Lightsheet 7, ZEISS), RPE1 cells were seeded onto 4-well cover glass chambered dishes (Lab-Tek; 1064716) or 4-well glass bottom ibidi^TM^ dish, and transferred to Leibovitz’s L15 medium (Invitrogen: 11415064) for imaging. For live-cell studies, 20,000 cells were seeded in each well 24 hours before imaging. Imaging was performed at 37°C using a full-stage incubation chamber setup to allow normal mitosis progression and cell cycle dynamics.

**FRAP & *di*SIM (Elyra7-RappOpto) imaging** For super-resolution live-cell imaging and FRAP, using Elyra 7 (ZEISS, Jena, Germany) equipped with UGA-42 Firefly (Rapp OptoElectronic GmbH, Germany), cells were seeded onto a 4-well ibidi^TM^ glass-bottom dish (ibidi^TM^; 80427) 24 hours and then changed to Leibovitz’s L15 medium (Invitrogen; 11415064) before imaging. Imaging was performed at 37°C using a full-stage incubation chamber, with 5s time interval, 3 Z planes, 0.5μm apart, in leap mode (9 Z-slice SIM), and acquired using a 63×/ 1.4 oil immersion objective. For photobleaching studies, a 473 nm laser was used at 100% power for 0.5 seconds. The Elyra 7 features Lattice SIM (Structured Illumination Microscopy) equipped with a *di*SIM image reconstruction algorithm, allowing fast and gentle super-resolution imaging (resolution of ∼60 nm in xy and a leap mode of accelerated volume imaging). Images were acquired with a 16-bit 512× 512 pixel PCO.edge 4.2 sCMOS. 60nm DNA origami (GATTAquant GmbH, Alexa Fluor 488 Dye) was used to assess the resolution of the setup.

**Particle size analysis and displacement** Automated particle analysis to count foci numbers in each cell was performed using Arivis™ software (Arivis Vision4D 4.1.2, ZEISS). For this, one representative SIM-processed image was selected, and a pipeline was generated by setting an intensity threshold for the segmentation of particles; Particles were classified based on mean intensity using an object feature filter. Once the pipeline was generated using one image, all *di*SIM-processed movies were analysed with the same pipeline.

For semi-automated measurements of the pre-bleached 53BP1 foci particle size, we used ZEN™ software for *di*SIM processing and Z-stack maximum projection, then used the Draw spline contour from the graphics tool to measure the area of the particles. The displacement of 53BP1 foci particles was measured as absolute displacement between the foci centroids at 5 sec before bleaching and 40 sec after recovery. The morphology measurements of 53BP1 foci were conducted with FIJI(image J) Analyse Particles tool to measure the perimeter and the Convex Hull of the foci. The ratio between the convex hull perimeter and ROI foci perimeter is a measure of the sphericity or ‘spikiness’. If the ratio is close to 1, then the foci are almost spherical; the closer the ratio goes towards zero, the spikier the foci.

**Immunoblotting** was performed on proteins separated on 8% SDS-PAGE gels by transferring them overnight onto PVDF membranes. Lysates were generated by treating cells with a Benzonase lysis buffer consisting of 75 mM HEPES, 150 mM NaCl, 1.5mM EGTA, 10 mM MgCl2, 10% Glycerol, 0.1% NP-40 and Benzonase 90 U/ml. Membranes were incubated in primary antibodies against eGFP (Abcam, ab290; 1:1000) and probed using secondary antibodies labelled with infrared fluorescent dyes, which were imaged using an iBright 1500 imager.

**SIM and FRAP analysis** SIM acquisition and *di*SIM processing were performed using ZEN Black™ software. Images and Movies were prepared using ZEN Blue™ software. Additional analysis of FRAP intensity measurement was conducted on Fiji^43^, Microsoft Excel™, and graphs were plotted using GraphPad Prism 9™. The equation for FRAP normalised relative fluorescence intensity calculation are:

Step 1. Photobleaching Rate (r) = (Fc-Fb)/(Fc_0_-Fb)

Step 2. Recovery Rate of ROI (R) = (Fi-Fb)/(Fi_0_-Fb)

Step 3. Normalised Recovery Rate of ROI = R/r

Fi represents the fluorescence intensity region of interest, Fi_0_ represents the fluorescence intensity region of interest before bleaching, Fc represents Fluorescence intensity control, Fc_0_ represents Fluorescence intensity control before bleaching, Fb represents the Fluorescence intensity background, and ROI represents the region of interest.

**Statistical analysis** was performed in Graph Pad Prism 9™ using statistics: Non-linear fit, and one-phase association analysis to determine half-life recovery of fluorescence intensity per compact and amorphous foci in FRAP. Independent group samples unpaired t-test and nonparametric tests, i.e. two-tailed Mann-Whitney test were used to determine the significance of any differences observed. Two-way ANOVA was used to make comparisons among groups. (*** P≤ 0.001, * P≤ 0.05, ns P> 0.05).

**Image dataset** Representative raw movies are uploaded to Zenondo to provide a shared resource for future users (DOI 10.5281/zenodo.13941718).

### Limitations of the study

The FRAP-SR method described here works well for subcellular structures that are larger than 60 nm. For those scenarios where structures are smaller than 60 nm, users may need to choose higher-resolution live-imaging methods. While FRAP-SR offers powerful insight into the compartmentalisation of diffraction-limited structures, FRAP-SR may not be readily scalable to hundreds of precisely targeted imaging events in the absence of automated foci detection tools. 53BP1 is an abundant protein which helped our analysis of eGFP signals, and if users are working with less abundant proteins they may need brighter fluorophores such as mNeonGreen.

## Author contributions

VMD designed the study and drafted the manuscript. CW helped edit the manuscript with VMD. CW conducted all the FRAP, SIM and LLS7 imaging studies. CW analysed images and generated figures for all panels, except the below. JCMG set up the Arivis Pro foci segmentation framework to analyse foci numbers (Figure 4C, Supplementary Figure 3B), and generated Figure 6E. SD analysed G1 bodies and prepared Supplementary Fig 5. CW and MC jointly analysed Figures 2C, 3C, 4A and 5B. NS conducted immunoblotting studies to characterize RPE1 cell lines for 53BP1-eGFP or Mad1-eGFP expression. VMN standardised and performed aphidicolin treatment and washes for aphidicolin release (Figures 5 and 6). SD cultured RPE1 for FRAP studies in Figures 1-4. Statistical analysis were performed by JCMG and CW. All authors commented on the manuscript.

## Acknowledgements

We acknowledge funding support from BBSRC, UKRI (BBR01003X/1, BB/W002698/1, BB/V018310/1 and BBT017716/1 to VMD), MRC, UKRI (MR/X013847/1 to VMD), QMUL (SBC8DRA2 and SBC9DRA2 to VMD), QMUL-ZEISS joint PhD studentship to MC and CR UK (C28598/A9787 to VMD) and CONAHCyT scholarship to JCMG (CVU no. 1042679). We acknowledge Sam Court and Petra Ungerer for infrastructure maintenance support. We thank Christoforos Efstathiou from the Draviam group for detailed comments and constructive feedback.

## References

1. Draviam, V.M., Xie, S., and Sorger, P.K. (2004). Chromosome segregation and genomic stability. Curr. Opin. Genet. Dev. 14, 120–125.

2. Huyen, Y., Zgheib, O., Ditullio, R.A., Jr, Gorgoulis, V.G., Zacharatos, P., Petty, T.J., Sheston, E.A., Mellert, H.S., Stavridi, E.S., and Halazonetis, T.D. (2004). Methylated lysine 79 of histone H3 targets 53BP1 to DNA double-strand breaks. Nature 432, 406–411.

3. Zgheib, O., Pataky, K., Brugger, J., and Halazonetis, T.D. (2009). An oligomerized 53BP1 tudor domain suffices for recognition of DNA double-strand breaks. Mol. Cell. Biol. 29, 1050–1058.

4. Wilson, M.D., Benlekbir, S., Fradet-Turcotte, A., Sherker, A., Julien, J.-P., McEwan, A., Noordermeer, S.M., Sicheri, F., Rubinstein, J.L., and Durocher, D. (2016). The structural basis of modified nucleosome recognition by 53BP1. Nature 536, 100–103.

5. Fradet-Turcotte, A., Canny, M.D., Escribano-Díaz, C., Orthwein, A., Leung, C.C.Y., Huang, H., Landry, M.-C., Kitevski-LeBlanc, J., Noordermeer, S.M., Sicheri, F., et al. (2013). 53BP1 is a reader of the DNA-damage-induced H2A Lys 15 ubiquitin mark. Nature 499, 50–54.

6. Schultz, L.B., Chehab, N.H., Malikzay, A., and Halazonetis, T.D. (2000). p53 binding protein 1 (53BP1) is an early participant in the cellular response to DNA double-strand breaks. J. Cell Biol. 151, 1381–1390.

7. Wang, B., Matsuoka, S., Carpenter, P.B., and Elledge, S.J. (2002). 53BP1, a mediator of the DNA damage checkpoint. Science 298, 1435–1438.

8. Hart, M., Adams, S.D., and Draviam, V.M. (2021). Multinucleation associated DNA damage blocks proliferation in p53-compromised cells. Commun Biol 4, 451.

9. Houtsmuller, A.B., Rademakers, S., Nigg, A.L., Hoogstraten, D., Hoeijmakers, J.H., and Vermeulen, W. (1999). Action of DNA repair endonuclease ERCC1/XPF in living cells. Science 284, 958–961.

10. Kilic, S., Lezaja, A., Gatti, M., Bianco, E., Michelena, J., Imhof, R., and Altmeyer, M. (2019). Phase separation of 53BP1 determines liquid-like behavior of DNA repair compartments. EMBO J. 38, e101379.

11. Zhang, L., Geng, X., Wang, F., Tang, J., Ichida, Y., Sharma, A., Jin, S., Chen, M., Tang, M., Pozo, F.M., et al. (2022). 53BP1 regulates heterochromatin through liquid phase separation. Nat. Commun. 13, 360.

12. Bothmer, A., Robbiani, D., Feldhahn, N., Gazumyan, A., Nussenzweig, A., and Nussenzweig, M. (2010). 53BP1 regulates DNA resection and the choice between classical and alternative end joining during class switch recombination. J. Exp. Med. 207, 855–865.

13. Difilippantonio, S., Gapud, E., Wong, N., Huang, C.-Y., Mahowald, G., Chen, H.T., Kruhlak, M.J., Callen, E., Livak, F., Nussenzweig, M.C., et al. (2008). 53BP1 facilitates long-range DNA end-joining during V(D)J recombination. Nature 456, 529–533.

14. Lukas, C., Savic, V., Bekker-Jensen, S., Doil, C., Neumann, B., Pedersen, R.S., Grøfte, M., Chan, K.L., Hickson, I.D., Bartek, J., et al. (2011). 53BP1 nuclear bodies form around DNA lesions generated by mitotic transmission of chromosomes under replication stress. Nat. Cell Biol. 13, 243–253.

15. Harrigan, J.A., Belotserkovskaya, R., Coates, J., Dimitrova, D.S., Polo, S.E., Bradshaw, C.R., Fraser, P., and Jackson, S.P. (2011). Replication stress induces 53BP1-containing OPT domains in G1 cells. J. Cell Biol. 193, 97–108.

16. Heemskerk, T., van de Kamp, G., Essers, J., Kanaar, R., and Paul, M.W. (2023). Multi-scale cellular imaging of DNA double strand break repair. DNA Repair 131, 103570.

17. Chapman, J.R., Sossick, A.J., Boulton, S.J., and Jackson, S.P. (2012). BRCA1-associated exclusion of 53BP1 from DNA damage sites underlies temporal control of DNA repair. J. Cell Sci. 125, 3529–3534.

18. Depes, D., Lee, J.-H., Bobkova, E., Jezkova, L., Falkova, I., Bestvater, F., Pagacova, E., Kopecna, O., Zadneprianetc, M., Bacikova, A., et al. (2018). Single-molecule localization microscopy as a promising tool for γH2AX/53BP1 foci exploration. Eur. Phys. J. D 72, 158.

19. Whelan, D.R., and Rothenberg, E. (2021). Super-resolution mapping of cellular double-strand break resection complexes during homologous recombination. Proc. Natl. Acad. Sci. U. S. A. 118. 10.1073/pnas.2021963118.

20. Mudumbi, K.C., Czapiewski, R., Ruba, A., Junod, S.L., Li, Y., Luo, W., Ngo, C., Ospina, V., Schirmer, E.C., and Yang, W. (2020). Nucleoplasmic signals promote directed transmembrane protein import simultaneously via multiple channels of nuclear pores. Nat. Commun. 11, 2184.

21. Mudumbi, K.C., Schirmer, E.C., and Yang, W. (2016). Single-point single-molecule FRAP distinguishes inner and outer nuclear membrane protein distribution. Nat. Commun. 7, 12562.

22. Löschberger, A., Novikau, Y., Netz, R., Spindler, M.-C., Benavente, R., Klein, T., Sauer, M., and Kleppe, I. (2021). Super-Resolution Imaging by Dual Iterative Structured Illumination Microscopy. bioRxiv, 2021.05.12.443720. 10.1101/2021.05.12.443720.

23. Guo, Y., Li, D., Zhang, S., Yang, Y., Liu, J.-J., Wang, X., Liu, C., Milkie, D.E., Moore, R.P., Tulu, U.S., et al. (2018). Visualizing Intracellular Organelle and Cytoskeletal Interactions at Nanoscale Resolution on Millisecond Timescales. Cell 175, 1430–1442.e17.

24. Chen, B.-C., Legant, W.R., Wang, K., Shao, L., Milkie, D.E., Davidson, M.W., Janetopoulos, C., Wu, X.S., Hammer, J.A., 3rd, Liu, Z., et al. (2014). Lattice light-sheet microscopy: imaging molecules to embryos at high spatiotemporal resolution. Science 346, 1257998.

25. Jullien, D., Vagnarelli, P., Earnshaw, W.C., and Adachi, Y. (2002). Kinetochore localisation of the DNA damage response component 53BP1 during mitosis. J. Cell Sci. 115, 71–79.

26. Lippincott-Schwartz, J., Snapp, E.L., and Phair, R.D. (2018). The development and enhancement of FRAP as a key tool for investigating protein dynamics. Biophys. J. 115, 1146–1155.

27. Pombo, A., Cuello, P., Schul, W., Yoon, J., Roeder, R.G., Cook, P.R., and Murphy, S. (1998). Regional and temporal specialization in the nucleus: a transcriptionally-active nuclear domain rich in PTF, Oct1 and PIKA antigens associates with specific chromosomes early in the cell cycle. EMBO J. 17, 1768–1778 – 1778.

28. Lottersberger, F., Karssemeijer, R.A., Dimitrova, N., and de Lange, T. (2015). 53BP1 and the LINC Complex Promote Microtubule-Dependent DSB Mobility and DNA Repair. Cell 163, 880–893.

29. Faustini, E., Panza, A., Longaretti, M., and Lottersberger, F. (2024). Quantitative analysis of nuclear deformations and DNA damage foci dynamics by live-cell imaging. Methods Cell Biol. 182, 247–263.

30. Dimitrova, N., Chen, Y.-C.M., Spector, D.L., and de Lange, T. (2008). 53BP1 promotes non-homologous end joining of telomeres by increasing chromatin mobility. Nature 456, 524–528.

31. Lezaja, A., Panagopoulos, A., Wen, Y., Carvalho, E., Imhof, R., and Altmeyer, M. (2021). RPA shields inherited DNA lesions for post-mitotic DNA synthesis. Nat. Commun. 12, 3827.

32. Bobkova, E., Depes, D., Lee, J.-H., Jezkova, L., Falkova, I., Pagacova, E., Kopecna, O., Zadneprianetc, M., Bacikova, A., Kulikova, E., et al. (2018). Recruitment of 53BP1 Proteins for DNA Repair and Persistence of Repair Clusters Differ for Cell Types as Detected by Single Molecule Localization Microscopy. Int. J. Mol. Sci. 19. 10.3390/ijms19123713.

33. Islam, A., Manjarrez-González, J.C., Song, X., Gore, T., and Draviam, V.M. (2024). Search for chromosomal instability aiding variants reveal naturally occurring kinetochore gene variants that perturb chromosome segregation. iScience 27, 109007.

34. Cuella-Martin, R., Hayward, S.B., Fan, X., Chen, X., Huang, J.-W., Taglialatela, A., Leuzzi, G., Zhao, J., Rabadan, R., Lu, C., et al. (2021). Functional interrogation of DNA damage response variants with base editing screens. Cell 184, 1081–1097.e19.

35. Löbrich, M., Rydberg, B., and Cooper, P.K. (1995). Repair of x-ray-induced DNA double-strand breaks in specific Not I restriction fragments in human fibroblasts: joining of correct and incorrect ends. Proc. Natl. Acad. Sci. U. S. A. 92, 12050–12054.

36. Núñez, M.I., Villalobos, M., Olea, N., Valenzuela, M.T., Pedraza, V., McMillan, T.J., and Ruiz de Almodóvar, J.M. (1995). Radiation-induced DNA double-strand break rejoining in human tumour cells. Br. J. Cancer 71, 311–316.

37. DiBiase, S.J., Zeng, Z.C., Chen, R., Hyslop, T., Curran, W.J., Jr, and Iliakis, G. (2000). DNA-dependent protein kinase stimulates an independently active, nonhomologous, end-joining apparatus. Cancer Res. 60, 1245–1253.

38. Obara, C.J., Nixon-Abell, J., Moore, A.S., Riccio, F., Hoffman, D.P., Shtengel, G., Xu, C.S., Schaefer, K., Pasolli, H.A., Masson, J.-B., et al. (2024). Motion of VAPB molecules reveals ER–mitochondria contact site subdomains. Nature 626, 169–176.

39. Sisario, D., Memmel, S., Doose, S., Neubauer, J., Zimmermann, H., Flentje, M., Djuzenova, C.S., Sauer, M., and Sukhorukov, V.L. (2018). Nanostructure of DNA repair foci revealed by superresolution microscopy. FASEB J., fj201701435.

40. Efstathiou, C., and Draviam, V.M. (2021). Electrically tunable lenses - eliminating mechanical axial movements during high-speed 3D live imaging. J. Cell Sci. 134. 10.1242/jcs.258650.

41. Lou, J., Priest, D.G., Solano, A., Kerjouan, A., and Hinde, E. (2020). Spatiotemporal dynamics of 53BP1 dimer recruitment to a DNA double strand break. Nat. Commun. 11, 5776.

42. Bleiler, M., Cyr, A., Wright, D.L., and Giardina, C. (2023). Incorporation of 53BP1 into phase-separated bodies in cancer cells during aberrant mitosis. J. Cell Sci. 136. 10.1242/jcs.260027.

43. Schindelin, J., Arganda-Carreras, I., Frise, E., Kaynig, V., Longair, M., Pietzsch, T., Preibisch, S., Rueden, C., Saalfeld, S., Schmid, B., et al. (2012). Fiji: an open-source platform for biological-image analysis. Nat. Methods 9, 676–682.

