## Supplementary figures and images for "FRAP-in-SR: Fluorescence recovery in the Super-Resolution regime reveals subcompartments of 53BP1 foci"

### Supplementary figures

Supplementary Figure 1

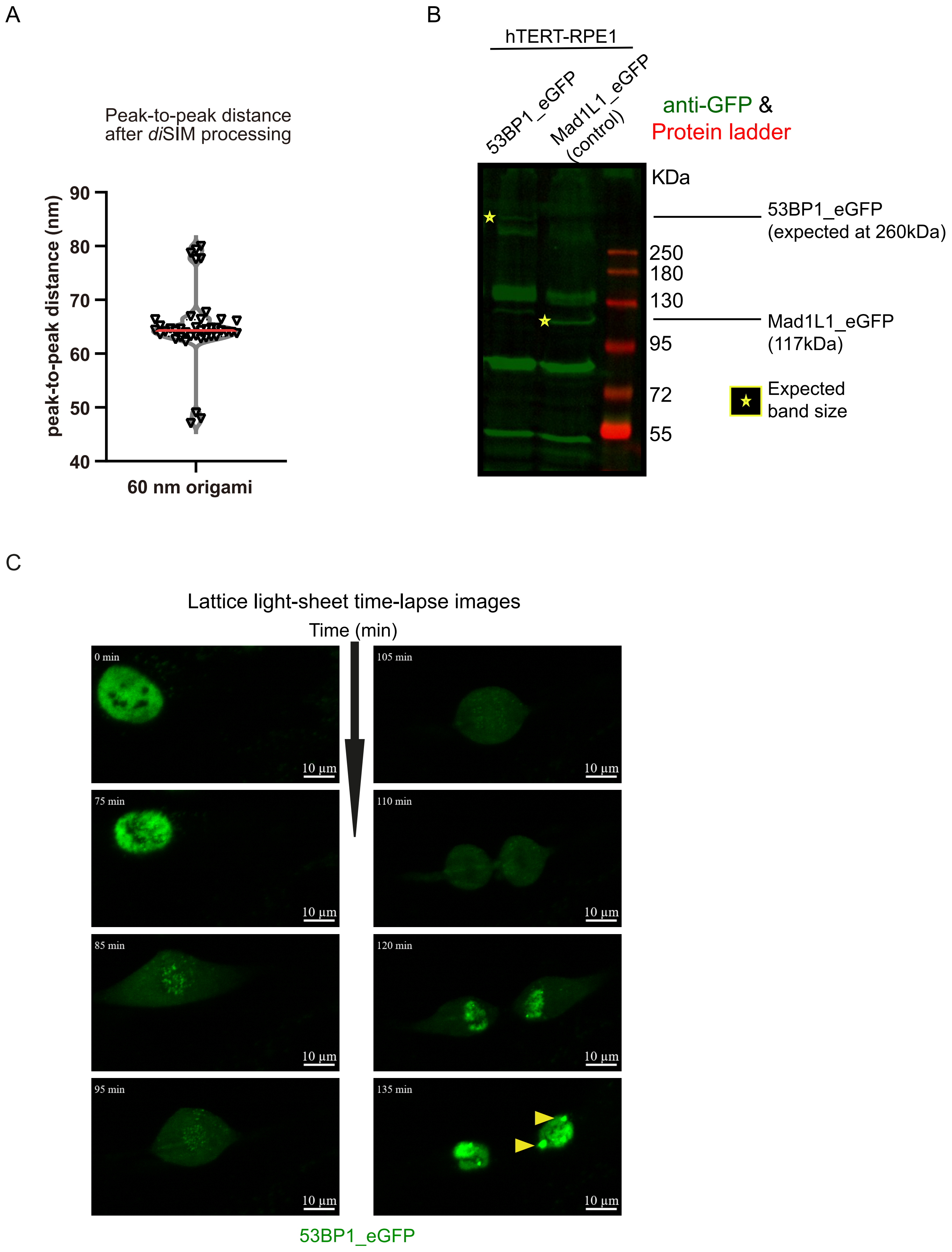

Supplementary figure 2

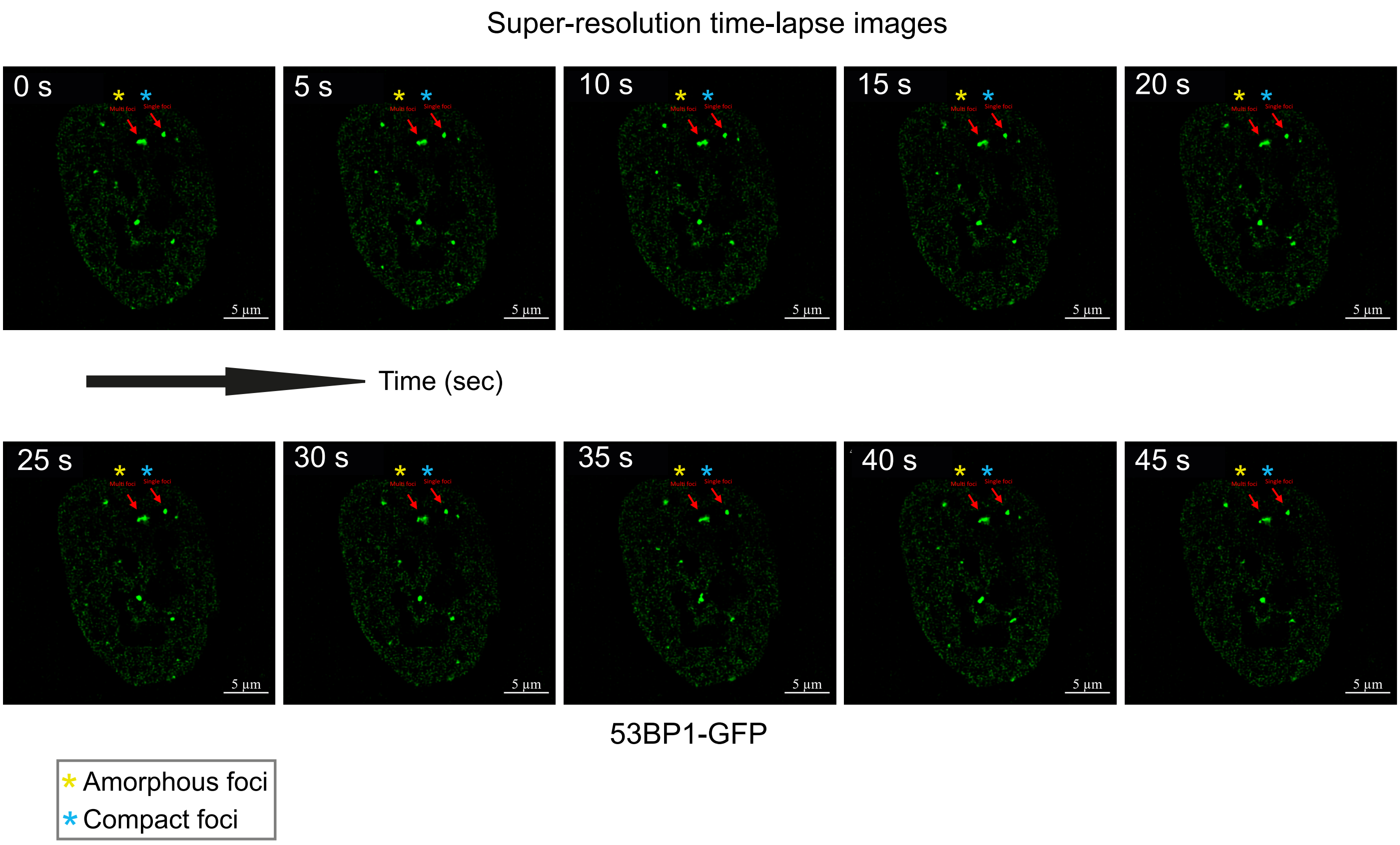

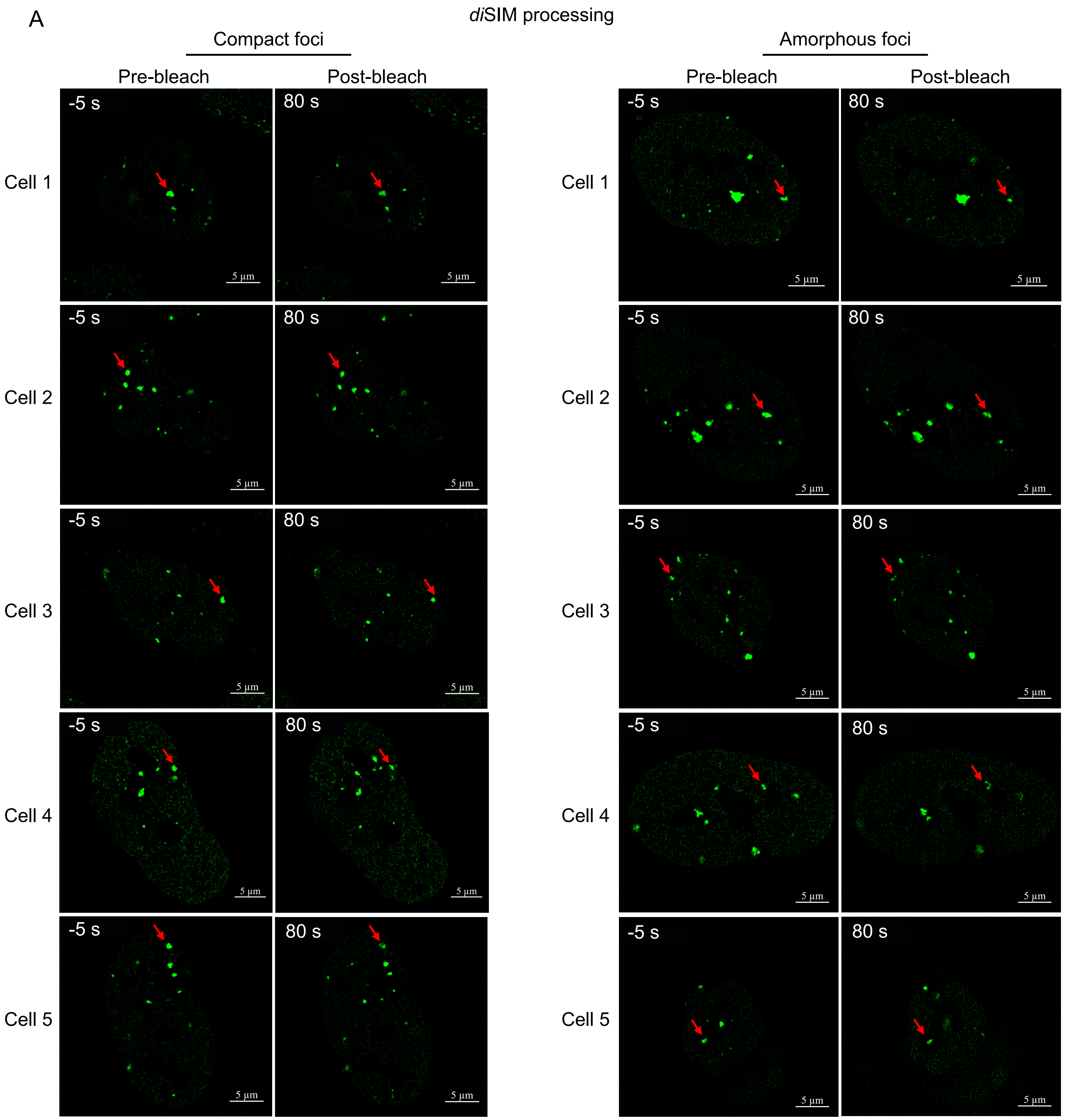

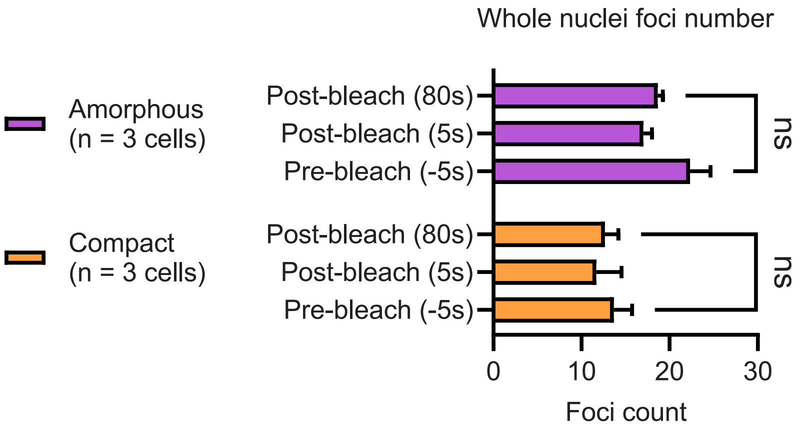

Supplementary Figure 4

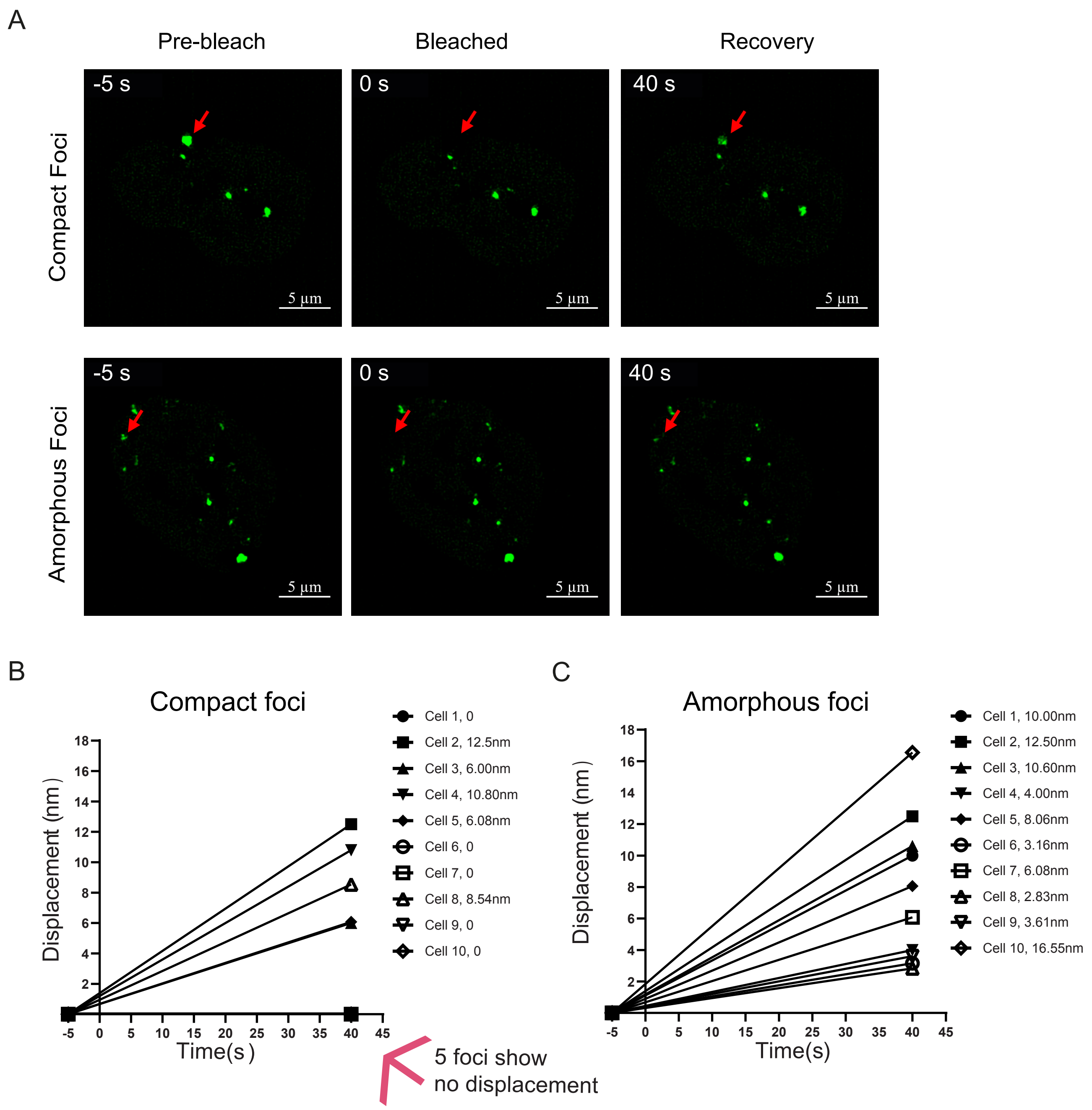

Supplementary Figure 5

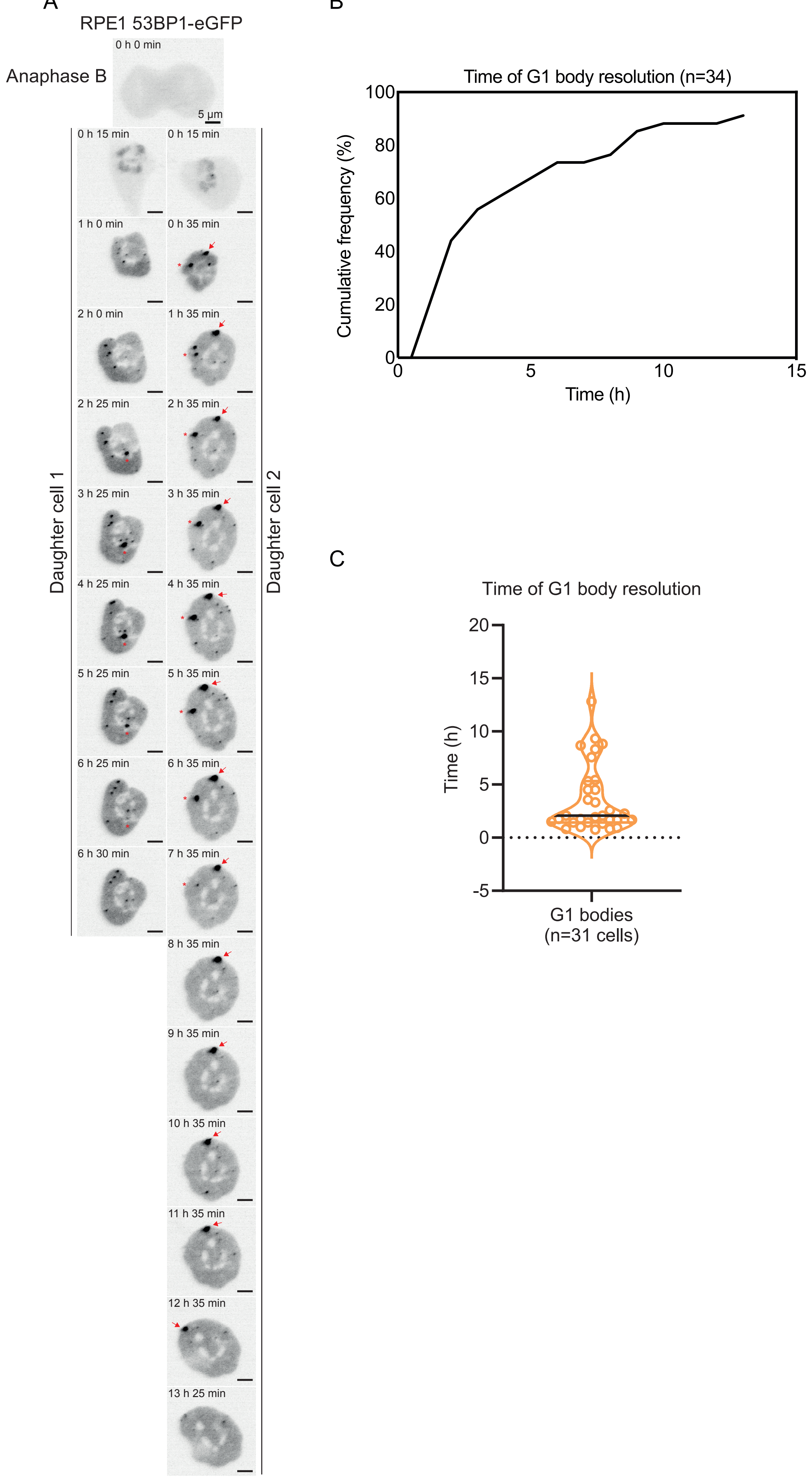

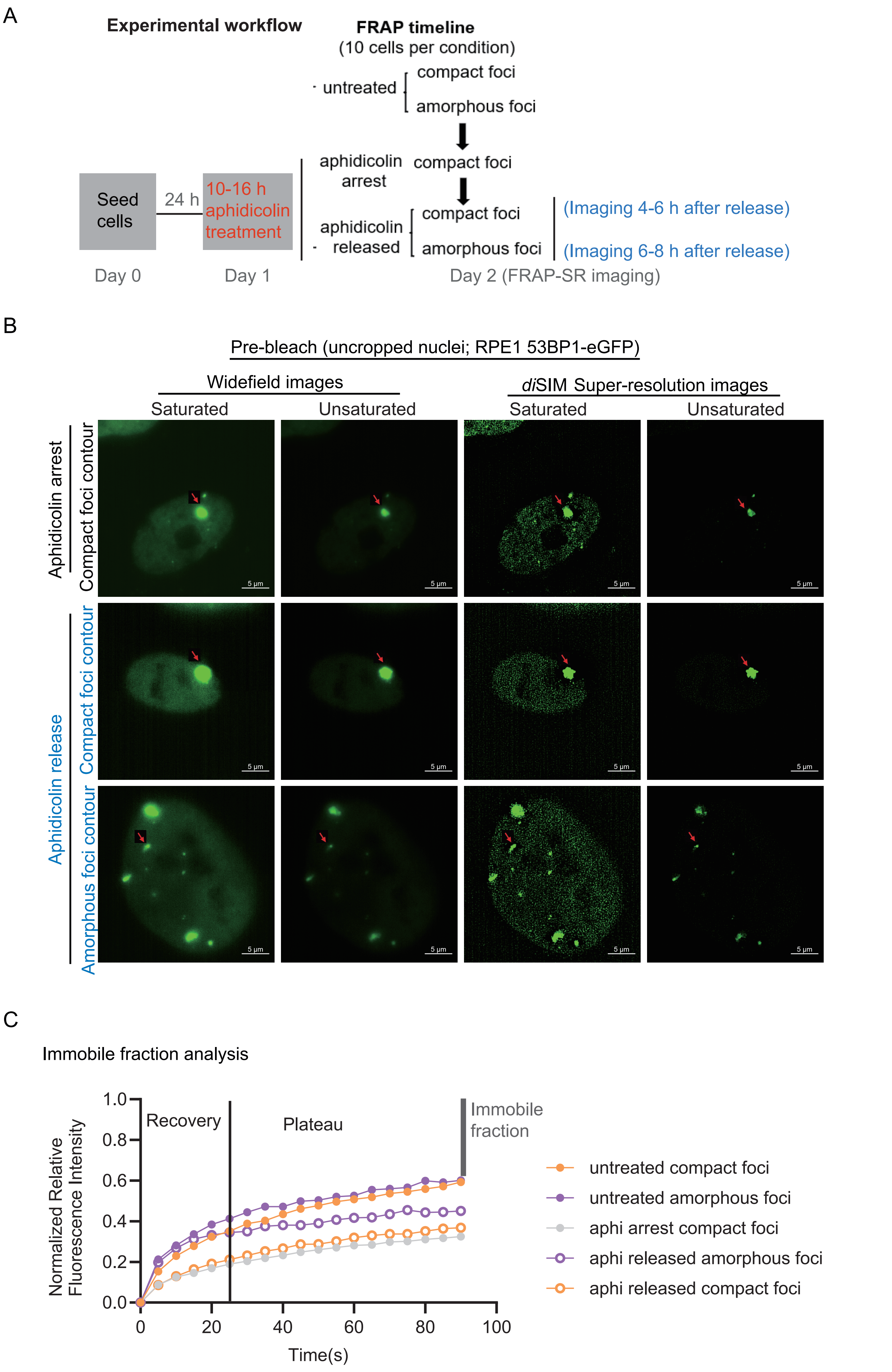
